# Reversible synaptic deficits in early-stage Batten disease

**DOI:** 10.1101/2025.11.12.687979

**Authors:** Masood Ahmad Wani, Chloe M. Hall, Thomas Mittmann, Benedikt Grünewald, Jakob von Engelhardt

## Abstract

Juvenile neuronal ceroid lipofuscinosis (JNCL, Batten Disease) is a childhood-onset, neurodegenerative, lysosomal storage disorder caused by mutations in the lysosomal gene *CLN3*. Progressive cognitive decline is a key clinical manifestation of JNCL, and no definitive treatment is currently available. The precise function of CLN3 in neurons remains unclear, hindering the development of targeted therapies. Using patch-clamp and AAV-mediated re-expression of *CLN3* in *Cln3*^Δex7/8^ mice, we demonstrate that CLN3 is essential for synaptic function. Loss of CLN3 caused defective synaptic vesicle release and reduced synaptic strength reflecting impairments in both pre- and postsynaptic function in *Cln3*^Δex7/8^ mice before the occurrence of neurodegeneration. Moreover, we observed reduced network bursting and deficits in intrinsic neuronal excitability, indicating early functional disturbances independent of storage burden and neuronal death in *Cln3*-deficient mice. To assess whether CLN3 is required for proper pre- and postsynaptic function, we re-expressed *CLN3* selectively in pre- and postsynaptic neurons. We found non-redundant requirements at both pre- and postsynaptic sites to sustain function. Importantly, AAV9-mediated gene rescue at early-disease stages corrected preexisting synaptic defects and restored function. Together, these findings demonstrate the requirement of CLN3 in maintaining synaptic function and show that the therapeutic window may extend to stages characterized by early functional deficits, highlighting the potential of targeted gene therapy to restore function and improve cognitive outcomes.

## Introduction

Juvenile neuronal ceroid lipofuscinosis (JNCL) is a childhood, neurodegenerative, lysosomal storage disorder (LSD) caused by mutations in the *CLN3* gene [30]. Children with JNCL experience progressive decline in cognition, accompanied by behavioral and psychological problems [25, 29]. The symptoms are thought to stem from neurodevelopmental and/or neurodegenerative processes. This view arises from observations of abnormal network activity and disruptions in lysosomal phospholipid catabolism early in the disease course, before symptoms and neurodegeneration occur [2, 46], whereas the severity of cognitive decline correlates with grey matter atrophy caused by neurodegeneration [15]. Together, these findings suggest that early disturbances in neurodevelopment may initiate neurodegenerative processes, leading to neuronal dysfunction and clinical symptoms. A distinct possibility is that synaptic dysfunction may drive cognitive impairment, rather than purely resulting from neurodegeneration. Synapses, as key components of neuronal circuits, are essential for learning and memory, and disruptions in their function can translate into cognitive impairments. Importantly, lysosomes, beyond their roles in degradation of macromolecules and cellular wastes, are increasingly recognized as modulators of synaptic homeostasis, influencing processes such as neurotransmitter release and plasticity [37]. Therefore, CLN3 deficiency may directly impair synaptic processes by disturbing endolysosomal function well before overt neuronal loss [10, 41, 42, 53, 54] and independent of neurodevelopmental processes.

We have previously identified such deficits in synaptic function in multiple brain regions including hippocampus at late-disease stage (14-month-old *Cln3*^-/-^ mice) [13]. However, recent studies have reported early network alterations that precede storage accumulation and neuronal loss in mice [2, 38]. Consistent with this, cortical organoid models of JNCL exhibit neurodevelopmental abnormalities and early changes in synaptic gene expression [12]. Whether these transcriptional abnormalities translate into neuronal and synaptic deficits that in turn drive network alterations remains unclear. Furthermore, it is still uncertain whether the synaptic changes observed in *Cln3*-deficient neurons represent irreversible developmental abnormalities or are directly caused by the absence of CLN3 in neurons and are therefore potentially reversible. This distinction is particularly important for defining the therapeutic window in which interventions could either prevent disease onset or reverse established symptoms. AAV-based gene therapies have shown safety and efficacy in pre-clinical models of JNCL. AAV-mediated *Cln3* rescue during the neonatal stages (P0-P2) and at one month of age had beneficial effects on the disease course and outcome at later stages in mice [1, 3, 20, 47, 57], supporting the promise of gene therapy for JNCL. However, none of the studies conducted to date have demonstrated a reversal of preexisting deficits.

To this end, we analyzed synaptic deficits in both young and adult mice and demonstrate that preexisting defects in presynaptic vesicle release and postsynaptic function can be reversed by selectively reintroducing *CLN3* into the pre- and postsynaptic neuron, respectively. These findings suggest a putative functional role for CLN3 at the synapse and indicate that treatment can restore synaptic function. Furthermore, we show that impairments in neuronal excitability and synaptic transmission precede substantial storage accumulation and cell death in the timeline of disease progression [2, 5, 10, 22, 35, 41, 42, 53, 54], supporting the notion that CLN3 plays a direct role in maintaining neuronal function. This argues against a purely neurodegenerative or neurodevelopmental classification of JNCL and suggests that the therapeutic window may extend to stages when neuronal deficits and early symptoms are already present.

## Material and methods

### Ethical approval and experimental animals

All animal experiments were approved by the Rhineland-Palatinate State Office of Health (G21-1-059, G24-1-001) and performed in compliance with the German Animal Welfare Act and the ARRIVE guidelines [24]. All mice used in the experiments are *Cln3*^Δex7/8^ (Jackson Lab 004685) and maintained in-house at the Universitätsmedizin Mainz animal facility (TARC). The mice were bred via heterozygous mating and wild-type littermates were used as controls. Mice were kept in 12-hour light-dark cycle with food and water ad libitum. Both sexes were used for the experiments.

### Surgical procedures and AAV injection

Animals were prepared for surgery and virus injections as previously described [14]. Briefly, subcutaneous injections of buprenorphine (0.1 mg/kg) and carprofen (10 mg/kg) were administered 30 minutes prior to surgery for analgesia. Deep anaesthesia was induced with 4% isoflurane in an induction chamber, after which mice were head-fixed in a stereotaxic frame. Anaesthesia was maintained throughout the surgery via nose-cone inhalation of isoflurane at 2 - 2.5% (v/v). The scalp was injected with the local anaesthetic lidocaine (4 mg/kg) before a small incision was made to expose the skull, and a burr hole was drilled over the injection site.

For presynaptic rescue experiments, a 400 nL cocktail of AAV9-hSyn-h*CLN3*-IRES-EGFP (Packgene) and AAV9-hSyn-hChR2(H134R)-mCherry (Addgene # 26976-AAV9; deposited by Karl Deisseroth), containing 2.5 × 10 genome copies of each virus, was bilaterally injected into the entorhinal cortex (AP = - 5.2 mm; ML = ±3.4 mm; DV = - 3.5 mm). Presynaptic rescue control groups (WT and *Cln3*^Δex7/8^ mice) received injections of AAV9-hSyn-hChR2(H134R)-mCherry only. For postsynaptic rescue experiments, 250 nL of AAV9-hSyn-h*CLN3*-IRES-EGFP (2.5 × 10 GC) was bilaterally injected into the dentate gyrus (AP = - 3.5 mm; ML = ±2.7 mm; DV = - 2.8 mm) of *Cln3*^Δex7/8^ mice. After each injection, the glass micropipette was left in place for 3 minutes before being slowly withdrawn. The scalp was sutured, and animals were monitored for 1-hour post-surgery during recovery from anaesthesia on a heated cage surface and subsequently returned to their home cages. Patch-clamp experiments were performed 2 to 3 weeks after virus injections.

### Electrophysiology

#### Slice preparation

Acute hippocampal slices from young and adult mice were prepared as described previously [13, 23]. For adult animals (4 and 6 months), buprenorphine (0.1 mg/kg body weight) was administered for analgesia prior to anesthesia. Then, animals were perfused intracardially with ice-cold high-sucrose aCSF solution containing (in mM): 212 sucrose, 26 NaHCO_3_, 1.25 NaH_2_PO_4_, 3 KCl, 0.2 mM CaCl_2_, 7 MgCl_2_, and 10 glucose. Horizontal hippocampal slices of 250 µm thickness were cut using a vibratome (Leica VT1200, Leica Microsystems, Germany). The slices were first recovered at 34°C in choline based aCSF containing (in mM): 87 NaCl, 2.5 KCl, 37.5 choline chloride, 25 NaHCO_3_, 1.25 NaH_2_PO_4_, 25 glucose, 0.5 CaCl_2_ and 7 MgCl_2_) for 20 minutes, then in standard aCSF containing (in mM): 125 NaCl, 25 NaHCO_3_, 1.25 NaH_2_PO_4_, 2.5 KCl, 2 CaCl_2_, 1 MgCl_2_, and 25 glucose for 10 minutes. Subsequently, slices were kept at room temperature until use.

#### Whole-cell patch clamp

All recordings were performed at 32°C in standard aCSF. Drug concentrations, when included, were 10 µM Gabazine (SR 95531 hydrobromide), 50 µM D-AP5, 10 µM CNQX and 1 µM Tetrodotoxin. Patch pipettes were pulled with a resistance of 3-5 MΩ from borosilicate glass capillaries using Sutter P-1000. Patch pipettes were filled with the following intracellular solutions depending on the experiment: (a) mEPSCs and intrinsic properties: (in mM) 130 potassium gluconate, 10 HEPES, 10 phosphocreatine-Na, 10 Na-gluconate, 0.3 GTP, 4 Mg² - ATP, and 4 NaCl (pH 7.2, adjusted with KOH). (b) mIPSCs: (in mM) 145 CsCl, 10 HEPES, 0.1 EGTA, 0.3 GTP, 2 Mg² -ATP, and 2 MgCl (pH 7.2, adjusted with CsOH). (c) AMPA/NMDA ratio and readily releasable pool (RRP): (in mM) 120 cesium gluconate, 10 CsCl, 8 NaCl, 10 HEPES, 10 phosphocreatine-Na, 0.3 GTP, 2 Mg² -ATP, and 0.2 EGTA (pH 7.3, adjusted with CsOH). (d) LTP recordings: (in mM) 120 cesium methanesulfonate, 10 CsCl, 8 NaCl, 10 HEPES, 10 phosphocreatine-Na, 0.3 GTP, 2 Mg² -ATP, and 0.2 EGTA (pH 7.3, adjusted with CsOH). Series resistance was monitored by applying 5 ms hyperpolarizing voltage steps and cells with a series resistance higher than 30 MΩ or a change over 25% during the experiment were excluded from analysis. Liquid junction potentials were not corrected. Data were recorded using HEKA EPC10 amplifier and stored with PatchMaster software (HEKA, Germany).

mEPSCs were recorded in the presence of Gabazine, and mIPSCs in the presence of CNQX. D-AP5 and Tetrodotoxin were added to the aCSF for mEPSC and mIPSC recording. Synaptic events were detected in Clampfit software using template match algorithm.

AMPA and NMDA receptor-mediated currents were evoked by stimulating the medial perforant pathway (MPP) and recorded from dentate gyrus granule cells (DG-GCs) at a holding potential of -70 mV and +40 mV, respectively. NMDAR-mediated current amplitudes were measured 25 ms after the stimulus artefact. Data were analyzed using Igor Pro Neuromatic.

To estimate readily-releasable pool (RRP) size, release probability (P_r_), and replenishment rate, the MPP was stimulated at 30 Hz for 1 second. The recordings were performed in the presence of D-AP5 and Gabazine. RRP, P_r_, and recovery rate were calculated using the ‘SMN’ fit method [45]. The RRP replenishment rate was estimated by fitting a single-exponential fit to individual eEPSCs, evoked at increasing inter-stimulus intervals after the initial train.

Whole-cell long-term potentiation (LTP) was induced as previously described [23]. Briefly, the postsynaptic neuron was depolarized to 0 mV for 3 minutes, followed by 120 pulse stimulation (2 Hz) of the medial perforant pathway (MPP: induced) while holding the cell at 0 mV. As a control, EPSCs from lateral perforant pathway (LPP: non-induced) were recorded 500 ms after the MPP stimulation. EPSCs within 60 s windows were averaged and normalized to baseline responses recorded prior to induction. Potentiation was quantified over the first and last 10 minutes post LTP induction.

For optogenetic stimulation of ChR2-expressing perforant pathway fibers, blue light (470 nm) from a TTL-controlled LED (Thorlabs, M470F1) was delivered via an optical fiber (Thorlabs, M87L01) coupled to a fiber-optic cannula. mCherry-labeled projections and EGFP-positive cells were visualized using infrared-DIC and epifluorescence microscopy.

#### High-density Microelectrode Array (HD-MEA) recordings

Acute hippocampal slices (250 µm thick) were prepared from adult mice as described above. The hippocampus, along with adjacent cortical tissue, was microdissected from the horizontal slice. For extracellular recordings, slices were placed in the 3.8 x 3.8 mm² recording area of an Accura HD-MEA chip (3Brain AG, Switzerland) containing 4096 electrodes and left for 5 minutes prior to recording to ensure stable contact with the electrode array. The aCSF containing (in mM): 125 NaCl, 25 NaHCO_3_, 1.25 NaH_2_PO_4_, 5 KCl, 2 CaCl_2_, 1 MgCl_2_, and 25 glucose was continuously oxygenated and exchanged in the recording chamber, while maintained at 32°C. Spontaneous activity was recorded for 5 minutes at a sampling rate of 18 kHz and 20 Hz high-pass filtered. All analysis were performed within the BrainWave 5 software (3Brain AG, Switzerland). For spike analysis, the data were bandpass filtered (100 Hz - 3000 Hz) and spikes were detected using the Precise Timing Spike Detection (PTSD) algorithm. Spike sorting was performed using principal component analysis (PCA), allowing for up to 4 clusters per electrode. Burst events were defined as at least five consecutive spikes with an inter-spike interval (ISI) of ≤ 100 ms.

### Immunohistochemistry and spine counts

DG granule cells were filled with 0.1 - 0.5% biocytin (Sigma-Aldrich) dissolved in intercellular solution via patch pipette for 15 minutes. Slices were fixed overnight in 4% formaldehyde at 4°C and washed 3x with PBS the next day. For immunofluorescence, slices were permeabilized and blocked in PBS containing 0.2% Triton X-100 and 5% BSA for 2 h at room temperature, then incubated overnight at 4°C with streptavidin-Alexa 488 or 594 (1:1000 in PBS) on a shaker. Following PBS washes, slices were mounted with Prolong Gold (Invitrogen). Confocal images of dendritic morphology were acquired using Zeiss LSM 710 with a 40x/1.30 Oil DIC objective (voxel size 0.2076 × 0.2076 × 1.2 µm³, 4.8 pixels/µm). The Z-stack images were processed using maximum intensity projection in ImageJ (v1.54b), and dendrites were semi-automatically traced with NeuronStudio (v0.9.92). Traced dendrites were analyzed in ImageJ using the SNT plugin, and Sholl intersections were calculated at 10 µm radii.

Dendritic spines in the medial molecular layer of the DG were imaged using Leica Stellaris 8 FALCON with a 100x/1.4 oil immersion objective and 5x digital zoom (voxel size 0.0196 × 0.0196 × 0.0701 µm³, 1024 × 1024 pixels). Spines were counted using NeuronStudio’s spine classifier and categorized as mushroom, thin, or stubby based on head-to-neck ratio, neck length-to-head diameter ratio, and head diameter: Mushroom: neck ratio ≥ 1.10, head diameter ≥ 0.350 µm. Thin: neck ratio < 1.10, length-to-head ratio ≥ 2.50 µm. Stubby: not meeting mushroom or thin criteria or head diameter < 0.350 µm.

### Statistics

Data are presented as mean [95% CI] or median [IQR/P_25_ - P_75_]. Normality was assessed using the Shapiro-Wilk test. For two-group comparisons, unpaired Student’s *t*-test was used for normally distributed data, and the Mann-Whitney *U* test for non-normal data. Multiple group comparisons were performed with one-way ANOVA followed by Bonferroni’s post hoc test (normal distributions) or Kruskal-Wallis test with Dunn’s post hoc comparisons (non-normal distributions). Differences in LTP induction across time windows and genotypes were analyzed with two-way ANOVA with Sidak’s post hoc test. *p* values are reported in figures, with the statistical test specified in figure legends and in supplementary (Tables 1-8). Statistical analyses were done in GraphPad Prism (version 10.2.3), and *p* < 0.05 was considered significant.

## Results

### Impaired intrinsic properties of neurons in *Cln3*^Δex7/8^ mice

We performed high-density extracellular recordings in acute hippocampal slices to identify potential aberrant network activity in *Cln3*^Δex7/8^ mice. We observed reduced spontaneous bursting in the dentate gyrus (DG), with a subtle reduction in the mean firing rate (Supplementary Fig. S1). This is in line with previous reports of reduced hippocampal network activity following perforant pathway stimulation [2, 46].

To probe whether alterations of intrinsic neuronal properties underlie the observed changes in network activity, we performed whole-cell patch clamp recordings from DG granule cells (DG-GCs) in 4 month-old *Cln3*^Δex7/8^ mice. Analysis of active properties revealed fewer action potentials in response to incremental suprathreshold current injections (Fig. 1c; Mann-Whitney *U* = 224, p = 0.0057, FDR-adjusted q = 0.043). Furthermore, action potential kinetics were altered: AP half-width and rise time were increased in *Cln3*^Δex7/8^ compared to WT mice, most likely reflecting changes in the expression or kinetics of voltage gated ion channels (Supplementary Fig. S2b). Passive membrane properties did not differ between genotypes (Fig. 1a and Supplementary Fig. S2a). To determine whether alterations in intrinsic membrane properties occur before overt symptoms, we recorded also from 3 week-old (presymptomatic) mice. Similar to the 4 month data, passive properties were unchanged, but a subtle reduction in action potential firing rate was observed (Fig. 1d; Mann-Whitney *U* = 181.5, p = 0.0063, FDR-adjusted q = 0.063), indicating early functional deficits. Together, these findings indicate early abnormalities in the intrinsic excitability of neurons in *Cln3*^Δex7/8^ mice, likely contributing to impaired network activity.

**Fig. 1.**
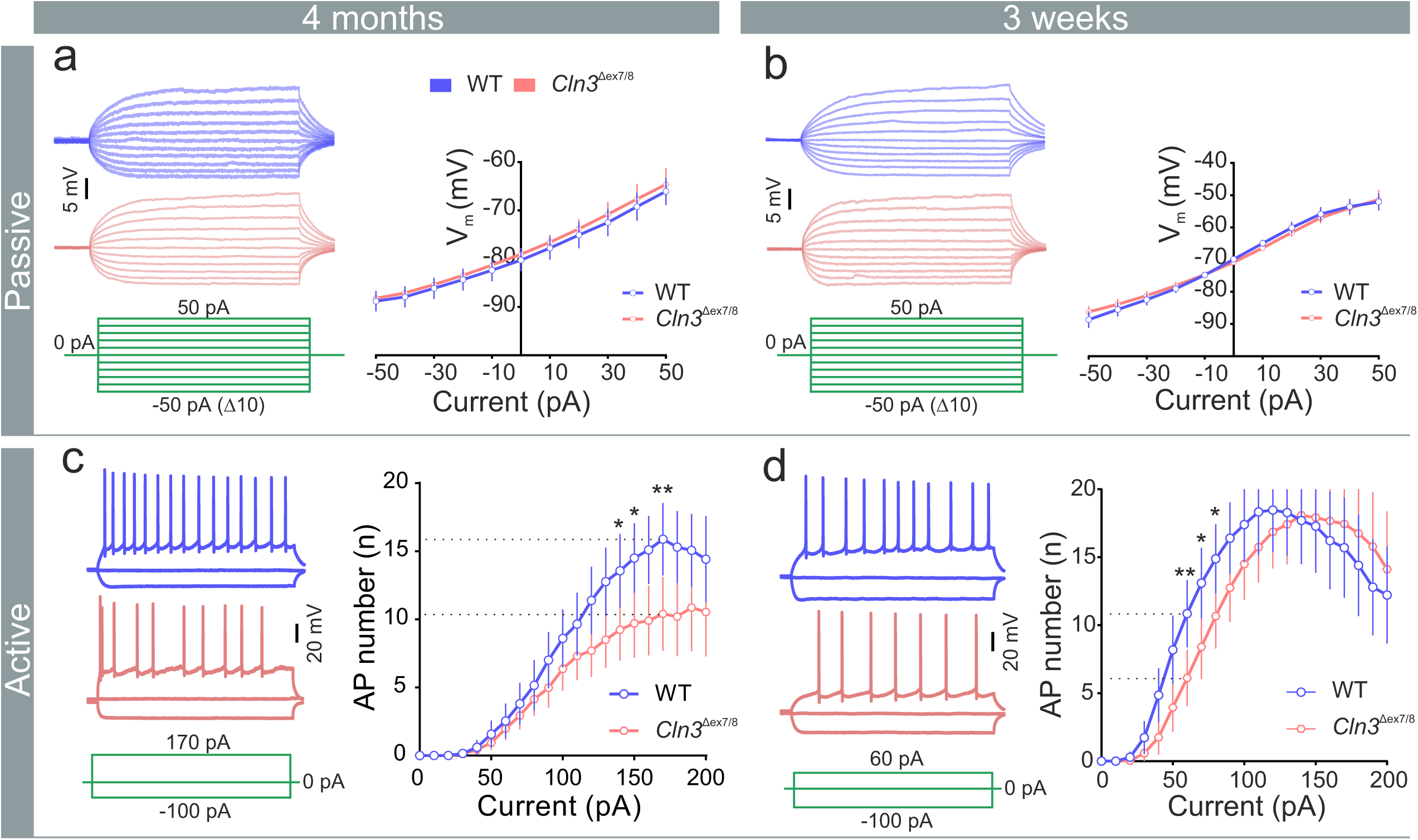
Subtle changes in intrinsic neuronal excitability in *Cln3*^Δex7/8^ mice. (a, b) Representative voltage responses from DG-GC to 300 ms current pulses: - 50 pA to 50 pA current steps in 10 pA increments and voltage-current relations (4-months: left, n= 20,32; N= 3,3 [WT, *Cln3*^Δex7/8^]; 3-weeks: right, n= 38,33; N= 5,4 [WT, *Cln3*^Δex7/8^]). (c, d) Firing pattern of DG-GCs in response to 1 second sub-threshold and hyperpolarizing current pulses: number of action potentials (AP) plotted against increasing current steps (4-months: left, n= 26,30; N= 4,5 [WT, *Cln3*^Δex7/8^]; 3-weeks: right, n= 27,24; N= 5,4 [WT, *Cln3*^Δex7/8^]). Data are mean ± SEM, **p < 0.01, *p < 0.05; Mann-Whitney test

### Excitatory synaptic transmission is impaired in *Cln3*^Δex7/8^ mice

To investigate whether synaptic dysfunction contributes to early network hypoactivity in hippocampus, we recorded miniature excitatory postsynaptic currents (mEPSCs) from 4-month-old mice. *Cln3*-deficient neurons exhibited a significant reduction in mEPSC frequency, with a subtle decrease in amplitude compared to WT, suggesting impairments in both pre- and postsynaptic mechanisms (4-months mEPSC frequency, median [P_25_ - P_75_]: WT, 1.96 [1.6 - 2.79] Hz vs *Cln3*^Δex7/8^, 0.75 [0.53 - 1.28] Hz and mEPSC amplitude, mean [95% CI]: WT, 13.61 [12.45 - 14.76] pA vs *Cln3*^Δex7/8^, 11.41 [10.48 - 12.34] pA) (Fig. 2a). A similar reduction in mEPSC frequency was also observed at 3-weeks of age, though without a change in amplitude, indicating a progressive decline in excitatory synaptic efficacy between 3 weeks and 4 months (3-weeks mEPSC frequency, median [P_25_ - P_75_]: WT, 3.16 [2.27 - 4.61] Hz vs *Cln3*^Δex7/8^, 2.11 [1.55 - 2.66] Hz and mEPSC amplitude, median [P_25_ - P_75_]: WT, 15.82 [13.57 - 17.36] pA vs *Cln3*^Δex7/8^, 15.56 [13.9 - 16.84] pA) (Fig. 2b). In contrast, miniature inhibitory postsynaptic currents (mIPSCs) showed no differences in frequency or amplitude at either age, suggesting that GABAergic transmission is largely preserved at presymptomatic disease stages (Fig. 2c and 2d). Taken together, these findings support our hypothesis that reduced synaptic drive may contribute to the hypoexcitable network phenotype observed in the DG at early stages and highlight a potential role for CLN3 in maintaining synaptic integrity during early disease progression.

**Fig. 2.**
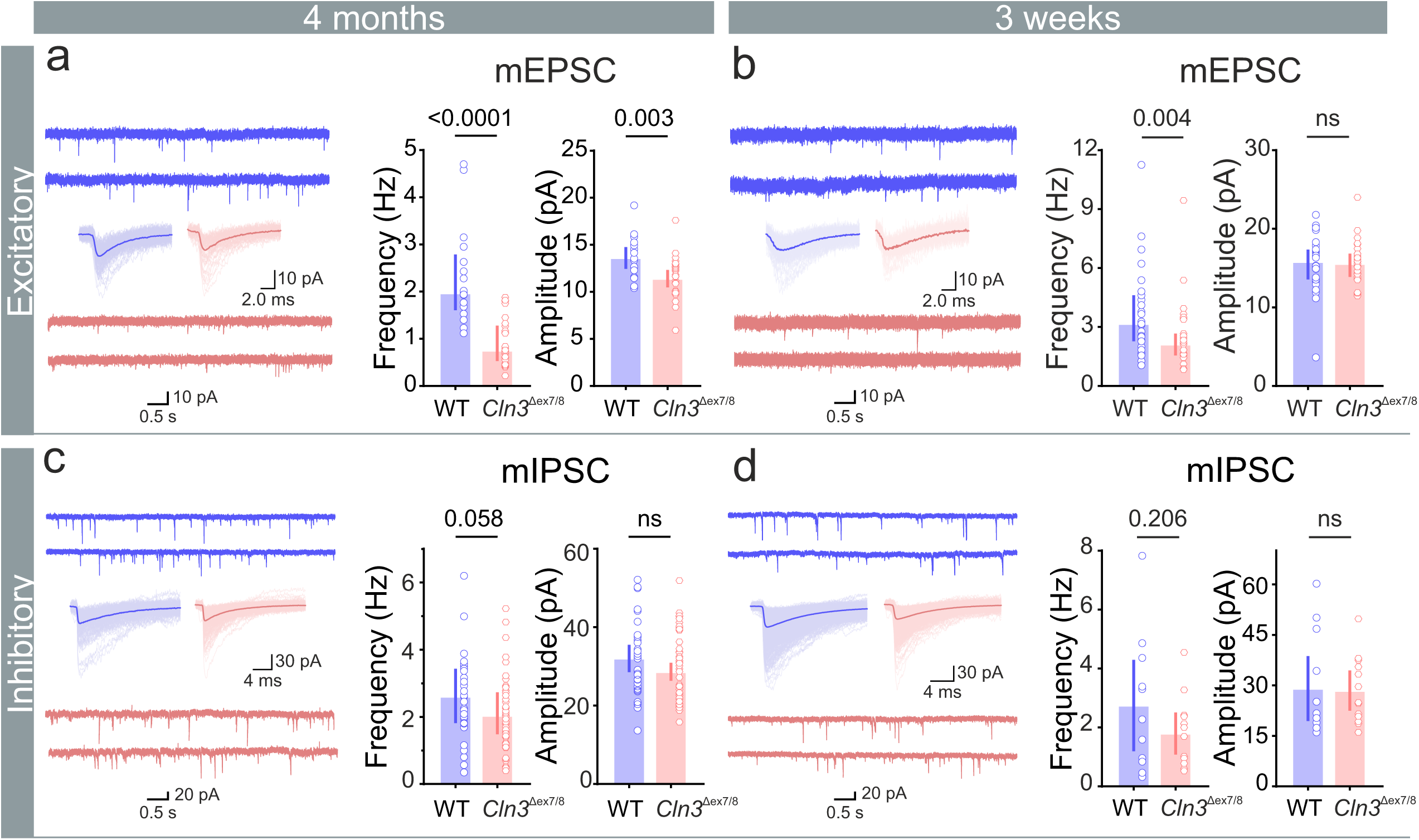
Impaired synaptic transmission in *Cln3*^Δex7/8^ mice. (a, b) Representative mEPSC traces from DG-GCs with summary bar plots of frequency and amplitude (4-months: left, n= 17,26; N= 3,3 [WT, *Cln3*^Δex7/8^]; 3-weeks: right, n= 28,31; N= 3,3 [WT, *Cln3*^Δex7/8^]). (c, d) Representative mIPSC traces from DG-GCs with summary bar plots of frequency and amplitude (4-months: left, n= 31,40; N= 5,5 [WT, *Cln3*^Δex7/8^]; 3-weeks: right, n= 12,13; N= 2,3 [WT, *Cln3*^Δex7/8^]). Data are median [IQR]; ns, non-significant; p-values from Mann-Whitney tests

### Synaptic strength is reduced in *Cln3*^Δex7/8^ mice

To determine whether loss of CLN3 affects synaptic strength, we stimulated the perforant pathway and recorded evoked AMPA and NMDA receptor-mediated currents from DG-GCs. The AMPA/NMDA ratio was significantly reduced in *Cln3*^Δex7/8^ mice compared to WT (A/N ratio, mean [95% CI]: WT, 1.86 [1.65 - 2.08] vs *Cln3*^Δex7/8^, 1.18 [1.01 - 1.35]) (Fig. 3b). To assess whether this reduced ratio was due to altered NMDA receptor function, we measured input-output responses of NMDAR-mediated currents across a range of stimulation intensities. We observed no difference in NMDAR-mediated current amplitudes between WT and *Cln3*^Δex7/8^ mice, suggesting that NMDA receptor density at these synapses is not affected by *Cln3* loss (Fig. 3c). Furthermore, the decay time constant (τ) of NMDAR-mediated currents was unaffected, indicating no major shifts in their subunit composition (Decay τ, mean [95% CI]: WT, 43.38 [37.50 - 49.26] ms vs *Cln3*^Δex7/8^, 44.61 [38.99 - 50.22] ms) (Fig. 3d). Together, these findings suggest that the reduced A/N ratio is likely due to a decrease in AMPAR numbers or channel conductivity. This is consistent with our observation of reduced mEPSC amplitude thus reflecting postsynaptic deficits in the absence of CLN3.

**Fig. 3.**
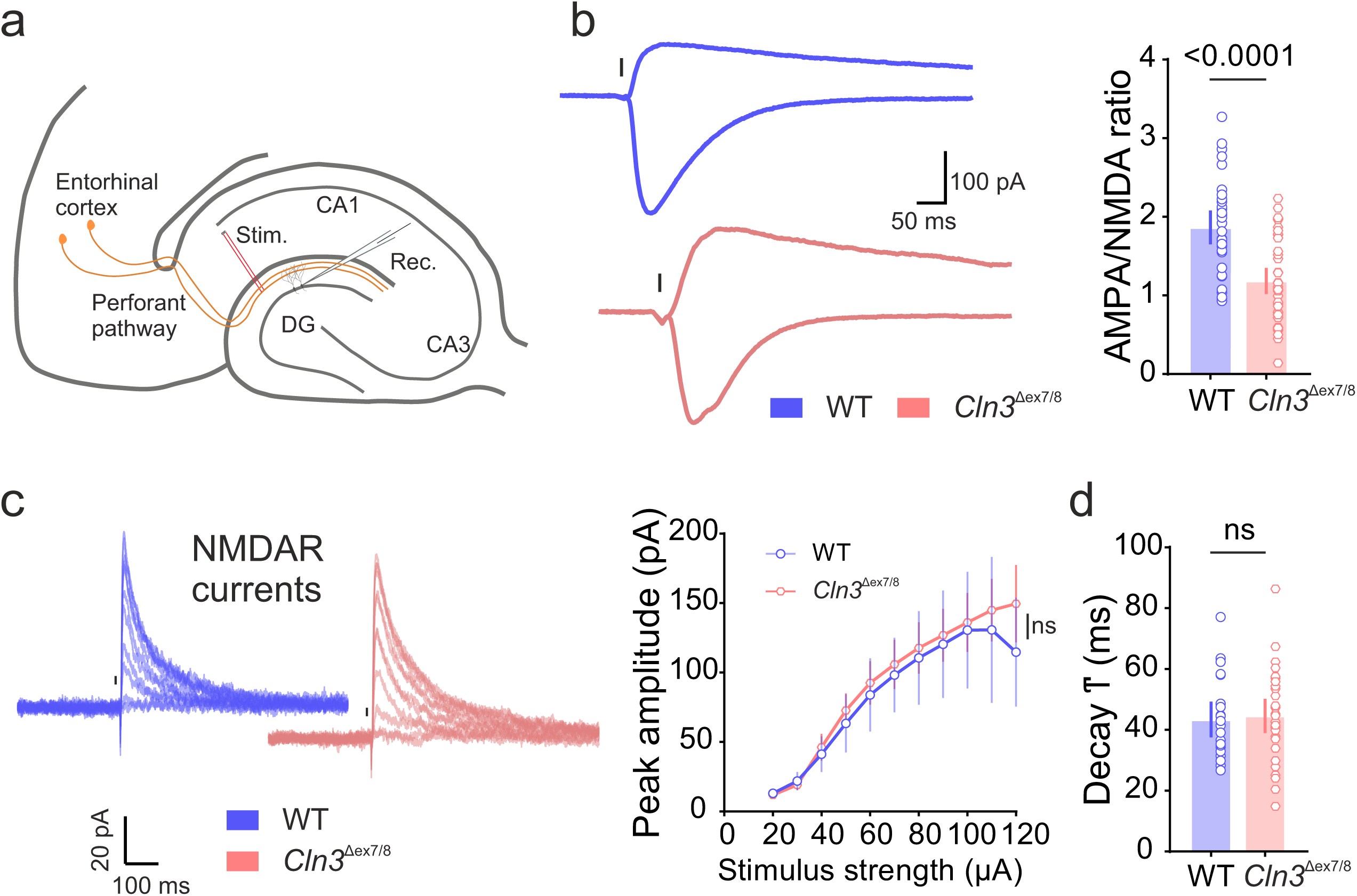
Loss of *Cln3* impairs synaptic strength in *Cln3*^Δex7/8^ mice. (a) Schematic of perforant pathway stimulation (Rec., recording pipette; Stim., stimulation electrode). (b) Example traces of evoked AMPA- and NMDA-receptor-mediated currents at -70 mV and +40 mV, respectively; stimulus artifacts removed (replaced by a black line) and summary bar plot of AMPA/NMDA ratio (n= 33,42; N= 5,6 [WT, *Cln3*^Δex7/8^]). (c) Representative traces of evoked NMDA receptor-mediated currents at +40 mV and peak amplitudes of NMDAR-mediated currents plotted against increasing stimulus intensities (μA). (d) Summary bar plot of decay time constant (τ) for NMDA receptor-mediated currents (n= 22,30; N= 2,2 [WT, *Cln3*^Δex7/8^]). Data are mean [95% CI]; ns, non-significant; p-values from Unpaired t-tests

### Neuronal spine number is reduced in *Cln3*^Δex7/8^ mice

Reduced mEPSC frequency may result from impaired synaptic vesicle release or from a decrease in the number of functional synapses, e.g., due to reduced dendritic spine density. To investigate this, DG-GCs were filled with biocytin via patch pipette, and spine density was quantified in the molecular layer of DG at 4-months. We observed a significant reduction in total dendritic spine density in granule cells of *Cln3*^Δex7/8^ mice compared to WT (spine count/10µm, mean [95% CI]: WT, 18.8 [17.9 - 19.7] vs *Cln3*^Δex7/8^, 16.05 [14.6 - 17.4]) (Fig. 4c). Further subclassification of spine types revealed a selective decrease in ‘mushroom’ and ‘thin’ spines, indicating a loss of mature spine subtypes (Fig. 4d). In addition, Sholl analysis revealed subtle alterations in dendritic arbour complexity in *Cln3*^Δex7/8^ mice compared to WT (Fig. 4e), suggesting that structural deficits may accompany synaptic impairment in the disease.

**Fig. 4.**
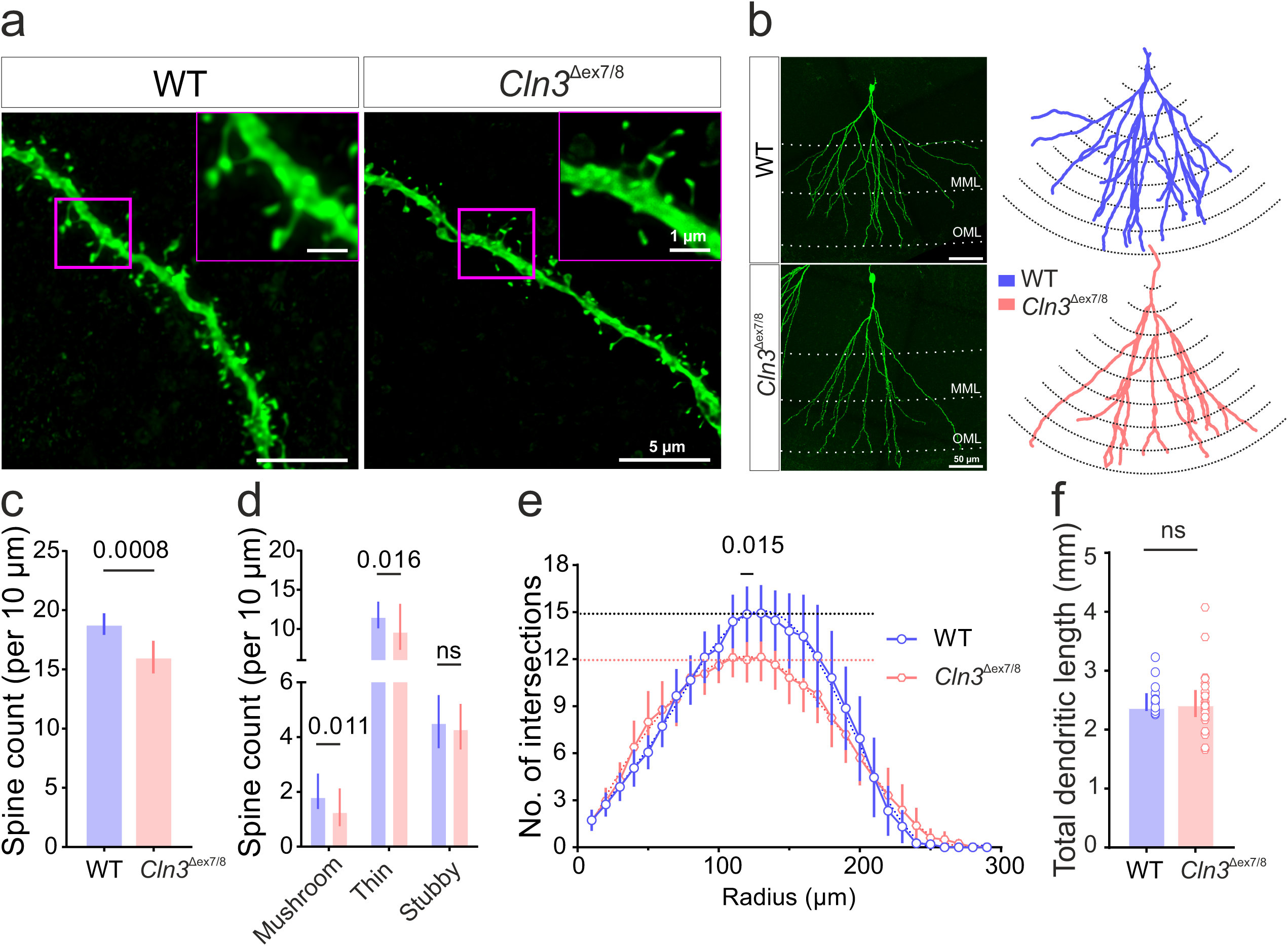
Hippocampal neurons of *Cln3*^Δex7/8^ mice have reduced spine number and dendritic complexity. (a) Confocal images of dendritic spines in the medial molecular layer of DG (inset scale bar, 1 μm). (b) Representative dendritic arbors of DG granule cells from WT and *Cln3*^Δex7/8^. Left: confocal images with dendrites visualized in green. Right: corresponding reconstructed tracings used for Sholl analysis. Scale bar: 50 μm. (c) Summary bar plots of total spine density and (d) spine subtype counts per 10 μm of dendritic length (n= 43,39; N= 8,5 [WT, *Cln3*^Δex7/8^]). (e) Sholl analysis quantifying dendritic intersections and (f) summary bar plot of total dendritic length (n= 17,26; N= 3,5 [WT, *Cln3*^Δex7/8^]). Data are mean [95% CI] in (c) and median [IQR] in (d, f); ns, non-significant; p-values from Unpaired t-test (total spine density) and Mann-Whitney tests with multiple-comparison FDR correction (spine subtype)

### Synaptic vesicle release probability (P_r_) is reduced and readily releasable pool size (RRP) is increased in *Cln3*^Δex7/8^ mice

The reduced mEPSC frequency suggests impaired presynaptic release of excitatory neurotransmitter vesicles in *Cln3*^Δex7/8^ mice. To assess how *Cln3* loss affects presynaptic vesicle dynamics, we estimated release probability (P_r_), readily releasable pool (RRP) size, and replenishment rate at perforant path-granule cell synapses by analyzing EPSCs evoked using a high frequency stimulation (HFS) protocol, as previously described [45]. We observed that *Cln3*^Δex7/8^ synapses exhibit a significantly reduced P_r,_ and an increased RRP size compared to WT (P_r_, mean [95% CI]: WT, 0.34 [0.31 - 0.38] vs *Cln3*^Δex7/8^, 0.27 [0.24 - 0.31] and RRP size, median [P_25_ - P_75_]: WT, 0.629 [0.498 - 0.782] nA vs *Cln3*^Δex7/8^, 0.815 [0.62 - 1.02] nA) (Fig. 5c). This suggests lower release probability and accumulation of synaptic vesicles in the absence CLN3. In contrast, the rate of recruitment of vesicles to the RRP during HFS (estimated from the slope of the cumulative EPSC plot) and replenishment rate after stimulation were similar between WT and *Cln3*^Δex7/8^ mice (Fig. 5d and Supplementary Fig. S5). This suggests that while SV release is impaired, the mobilization of vesicles from the reserve pool remains intact during high neuronal activity.

**Fig. 5.**
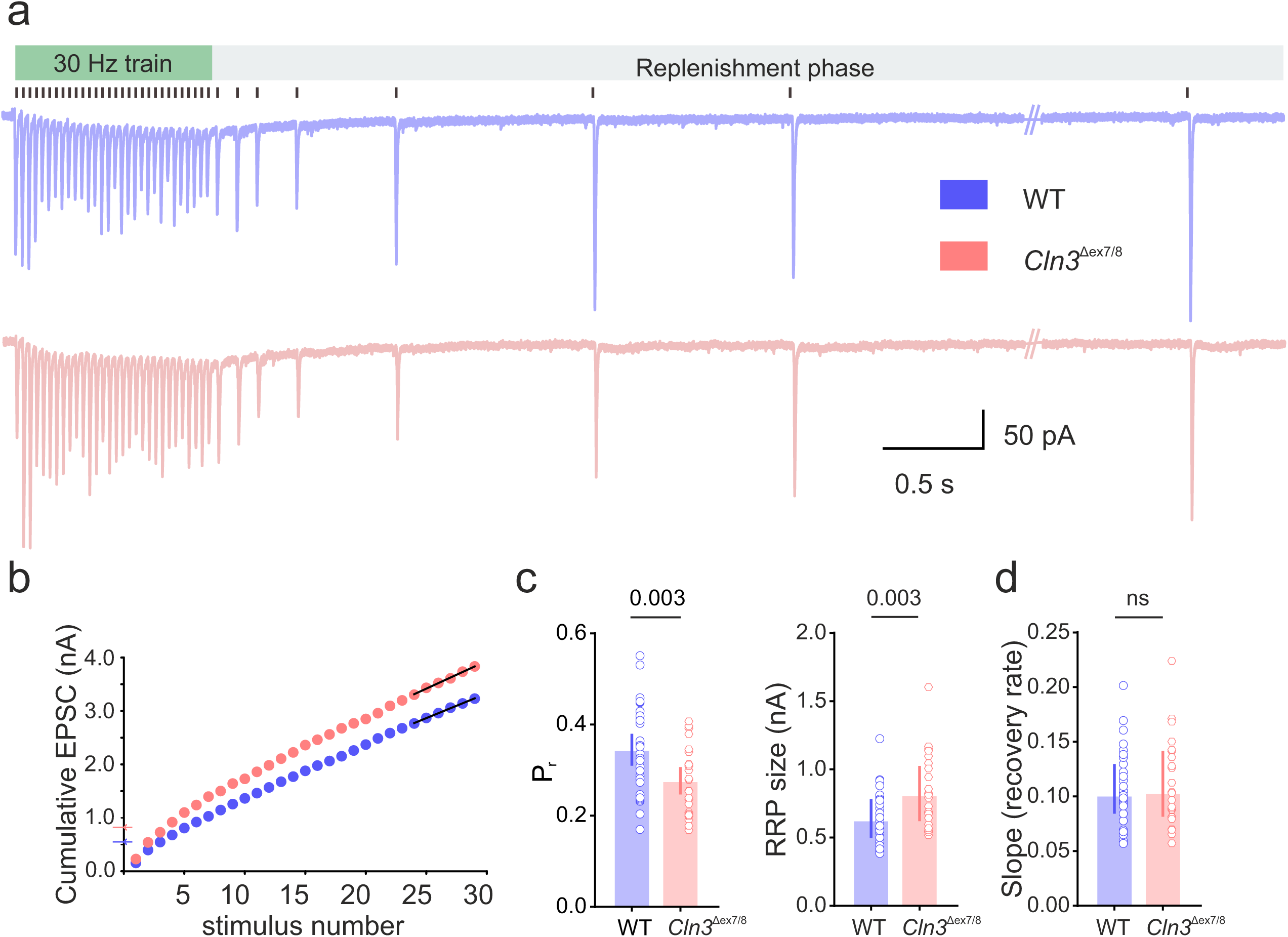
Synapses in *Cln3*^Δex7/8^ mice have reduced vesicle release probability. (a) Representative trace of evoked EPSCs in response to 30 Hz perforant pathway stimulation (black vertical lines above the traces indicate stimulus times). (b) Cumulative EPSC plot with linear fit (black line) of the last 6 points (steady state). (c) Summary bar plots of release probability (P_r_), readily releasable pool (RRP) size, and (d) slope (recovery rate) (n= 31,25; N= 4,3 [WT, *Cln3*^Δex7/8^]). Data are mean [95% CI] for P_r_ and median [IQR] for RRP size and slope; ns, non-significant; p-values from Unpaired t-test (P_r_) and Mann-Whitney tests (RRP size, slope)

### Long-term plasticity is intact in *Cln3*^Δex7/8^ mice

Synapses are key sites of neuronal plasticity potentially serving as substrates for memory formation and storage. Given that memory impairments are reported in both JNCL patients and animal models, we assessed long-term potentiation (LTP) in the hippocampus at 4-months (early disease stage). LTP was induced at the medial perforant path (MPP)-granule cell (GC) synapse, while the lateral perforant pathway (LPP) served as a control.

We examined synaptic strength during the first 10 minutes and the last 10 minutes following induction. Both WT and *Cln3*^Δex7/8^ mice showed comparable levels of potentiation at the MPP-GC synapse at both time points (first 10 min, mean [95% CI]: WT, 207.9 [162.1 - 253.8] pA vs *Cln3*^Δex7/8^, 183.4 [138 - 228.8] pA and last10 min, mean [95% CI]: WT, 181.1 [140.3 - 221.9] pA vs *Cln3*^Δex7/8^, 183.8 [129.6 - 238] pA (Fig. 6d). Likewise, the heterosynaptic LTD observed at the non-stimulated LPP was unaffected. These findings suggest that the core mechanisms underlying long-term synaptic plasticity remain intact at early stages despite early defects in synaptic transmission.

**Fig. 6.**
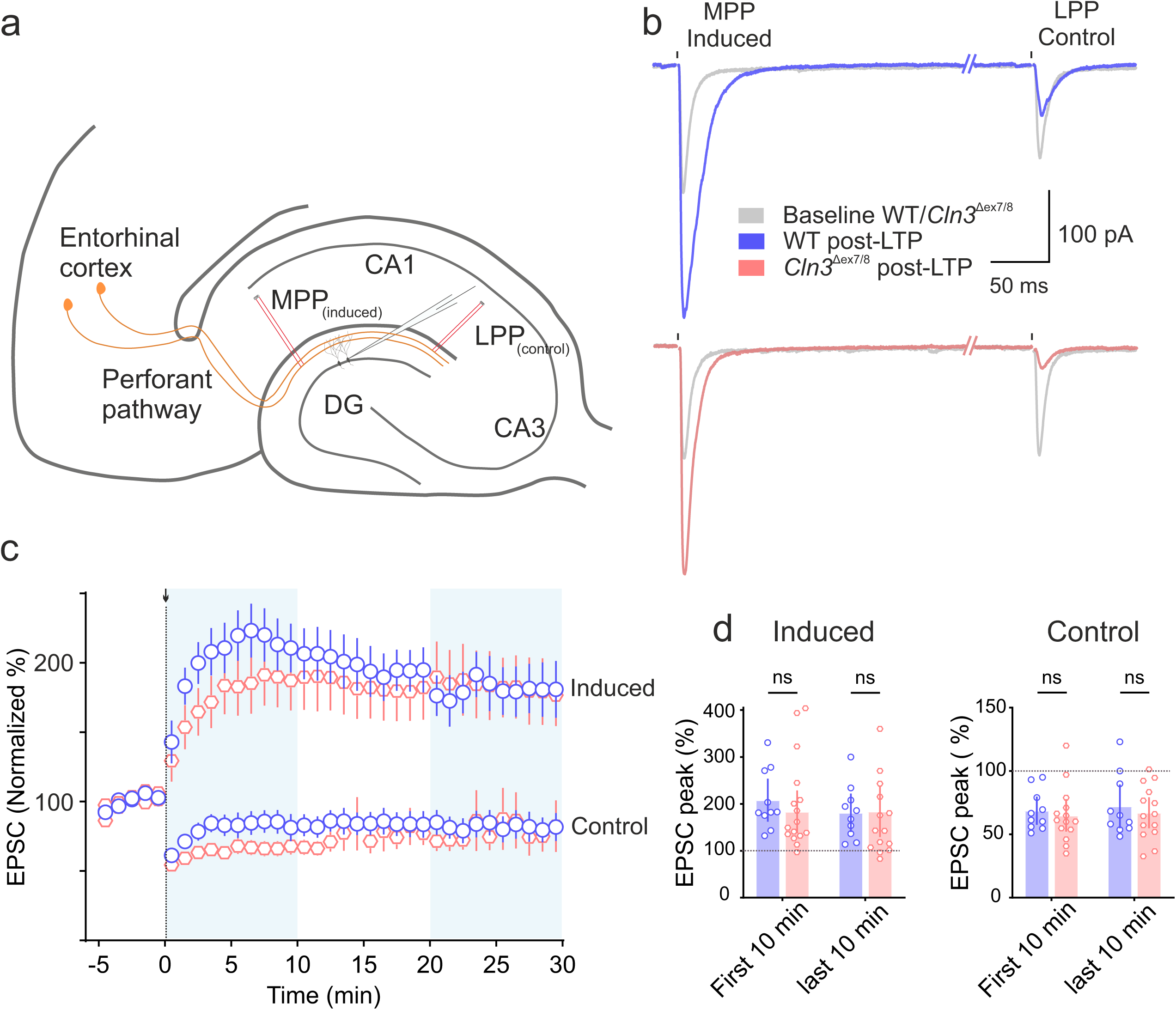
Hippocampal long-term potentiation (LTP) is intact in *Cln3*^Δex7/8^ mice. (a) Schematic of LTP induction in the DG via stimulation of the perforant pathway (MPP, medial perforant pathway, induced; LPP, lateral perforant pathway, control). (b) Example traces of evoked synaptic currents at baseline (grey) and after LTP induction for WT and *Cln3* (colored). (c) Time course of normalized EPSC peaks showing LTP induced at the MPP-GC synapse at time 0 (black arrow), while LPP synapses show heterosynaptic depression. Data are mean ± SEM. (d) Summary bar plots of induced and control EPSC peaks normalized to baseline for the first and last 10 minutes (n= 13,14; N= 6,5 [WT, *Cln3*^Δex7/8^]). Data are mean [95% CI]; ns, non-significant; two-way RM ANOVA with Sidak post hoc test

### Presynaptic re-expression of *CLN3* normalizes synaptic vesicle release

The loss of CLN3 disrupts both pre- and postsynaptic function. But whether these defects reflect independent roles for CLN3 in each site of the synapse or arise secondarily remains unclear. To address this question, we expressed *CLN3* selectively in either presynaptic or postsynaptic neurons and assessed the corresponding functional outcomes. For presynaptic targeting, we co-injected two rAAVs into the entorhinal cortex, one encoding *CLN3* and the other channelrhodopsin (ChR2). This approach enabled selective stimulation of perforant path fibers in the dentate gyrus arising from transgene-expressing neurons (Fig. 7a and b) The high overlap of green (CLN3) and red (ChR2) signal in the entorhinal cortex suggests that the majority of ChR2- positive neurons express CLN3. To estimate presynaptic vesicle release properties following rescue, perforant pathway axons expressing ChR2 were optically stimulated at 30 Hz to mimic high-frequency electrical stimulation and synaptic responses were recorded from the DG-GCs. Re-expression of *CLN3* in the presynaptic neurons for 3-weeks was sufficient to normalize P_r_ and RRP size to WT levels (P_r_, median [P_25_ - P_75_]: WT, 0.277 [0.240 - 0.396] vs *Cln3*^Δex7/8^, 0.186 [0.136 - 0.262] vs *Cln3*^Δex7/8^ + AAV9.*CLN3*, 0.257 [0.233 - 0.338] and RRP size, median [P_25_ - P_75_]: WT, 0.477 [0.352 - 0.702] nA vs *Cln3*^Δex7/8^, 0.920 [0.598 - 1.12] nA vs *Cln3*^Δex7/8^ + AAV9.*CLN3*, 0.632 [0.497 - 0.770] nA) (Fig. 7d). Similar to the experiments using electrical stimulation, the rate of recruitment of vesicles to the RRP (slope of cumulative EPSC plot) and the replenishment rate after optical stimulation were unaffected by CLN3 deficiency. Overexpression of *CLN3* did not change these parameters compared to WT (Fig. 7e and Supplementary Fig. S6). As there is no rAAV expression in DG granule cells (Fig. 7b), the observed changes in synapse function are due to re-expression of CLN3 in the presynaptic compartment. Our findings provide proof of concept that CLN3 plays a functional role within the presynaptic compartment and reveal that targeted neuronal rescue of CLN3 is sufficient to restore synaptic function.

**Fig. 7.**
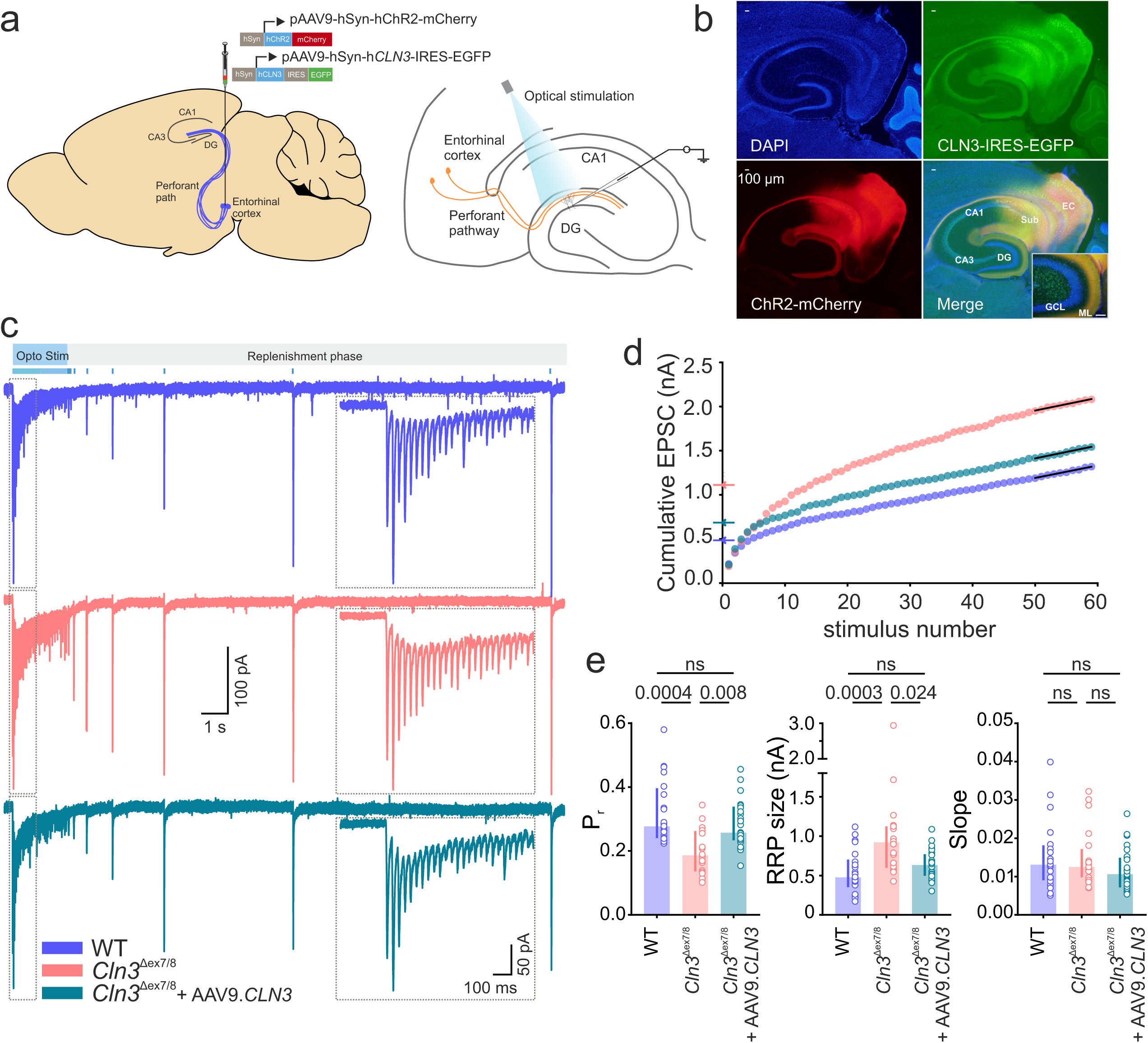
Re-expression of *CLN3* in the presynaptic neurons of *Cln3*^Δex7/8^ mice improves defective synaptic release. (a) Schematic of AAV injections into the entorhinal cortex (EC) to re-introduce *CLN3* in presynaptic neurons, combined with optogenetics for selective activation of *CLN3*-expressing perforant pathway axons. (b) Representative fluorescence images showing CLN3-EGFP and ChR2-mCherry expression in the EC and perforant pathway; merged image shows their colocalization in the molecular layer (ML) of DG (inset, scale bar: 100 µm). Sub, subiculum; GCL, granule cell layer. (c) Representative trace of evoked EPSCs in response to 30 Hz optical stimulation of the perforant pathway (blue vertical lines above the traces mark stimulus times). The eEPSC traces in the dotted box shows a zoomed-in portion of the high-frequency optical stimulation period. (d) Cumulative EPSC plot with linear fit (black line) of the last 10 points (steady state). (e) Summary bar plots of release probability (P_r_), readily releasable pool (RRP) size, and slope (n= 22,20,26; N= 3,3,4 [WT, *Cln3*^Δex7/8^, *Cln3*^Δex7/8^ + AAV9.*CLN3*]). Data are median [IQR]; ns, non-significant; p-values from Kruskal-Wallis test followed by Dunn’s multiple comparisons test

### Postsynaptic re-expression of *CLN3* normalizes synaptic strength

Next, we selectively re-expressed *CLN3* in DG granule cells by stereotactic rAAV injection into the DG to analyze the assumed role of CLN3 in the postsynaptic compartment (Fig. 8a and b). To identify the rAAV infected neurons during patch-clamp recordings, *CLN3* was expressed along with EGFP (Fig. 8c). EGFP-positive DG-GCs were patched and the perforant pathway was electrically stimulated to record AMPA- and NMDA-receptor-mediated currents. The AMPA/NMDA ratio in *CLN3* re-expressing neurons were higher than in neurons of *Cln3*^Δex7/8^ mice and comparable to the ratio in WT mice indicating normalization of synaptic strength in the rescued neurons (A/N ratio, median [P_25_ - P_75_]: *Cln3*^Δex7/8^, 1.43 [0.98 - 1.76] vs *Cln3*^Δex7/8^ + AAV9.*CLN3*, 1.73 [1.39 - 2.33]) (Fig. 8e). The rescue of A/N ratio following postsynaptic re-expression of *CLN3* in the *Cln3*^Δex7/8^ mice suggests that CLN3 may have unique roles at both the synaptic sites. We further recorded mEPSC from the postsynaptic rescued neurons and observed a rescue of mEPSC amplitude but not the frequency indicating that the re-expression of *CLN3* in postsynaptic compartment is insufficient to correct presynaptic defects (mEPSC frequency, median [P_25_ - P_75_]: *Cln3*^Δex7/8^, 0.40 [0.18 - 0.74] Hz vs *Cln3*^Δex7/8^ + AAV9.*CLN3*, 0.34 [0.27 - 0.51] Hz and mEPSC amplitude, mean [95% CI]: *Cln3*^Δex7/8^, 12.42 [11.84 - 13.01] pA vs *Cln3*^Δex7/8^ + AAV9.*CLN3*, 13.82 [12.62 - 15.01] pA) (Fig. 8f). To further confirm this, we measured RRP and the P_r_, two presynaptic metrics changed in the *Cln3*^Δex7/8^ mice, from the postsynaptic rescued neurons and found no change in both (Supplementary Fig. S7). These findings highlight the requirement of CLN3 in both the synaptic compartments to fully salvage synaptic function in the disease.

**Fig. 8.**
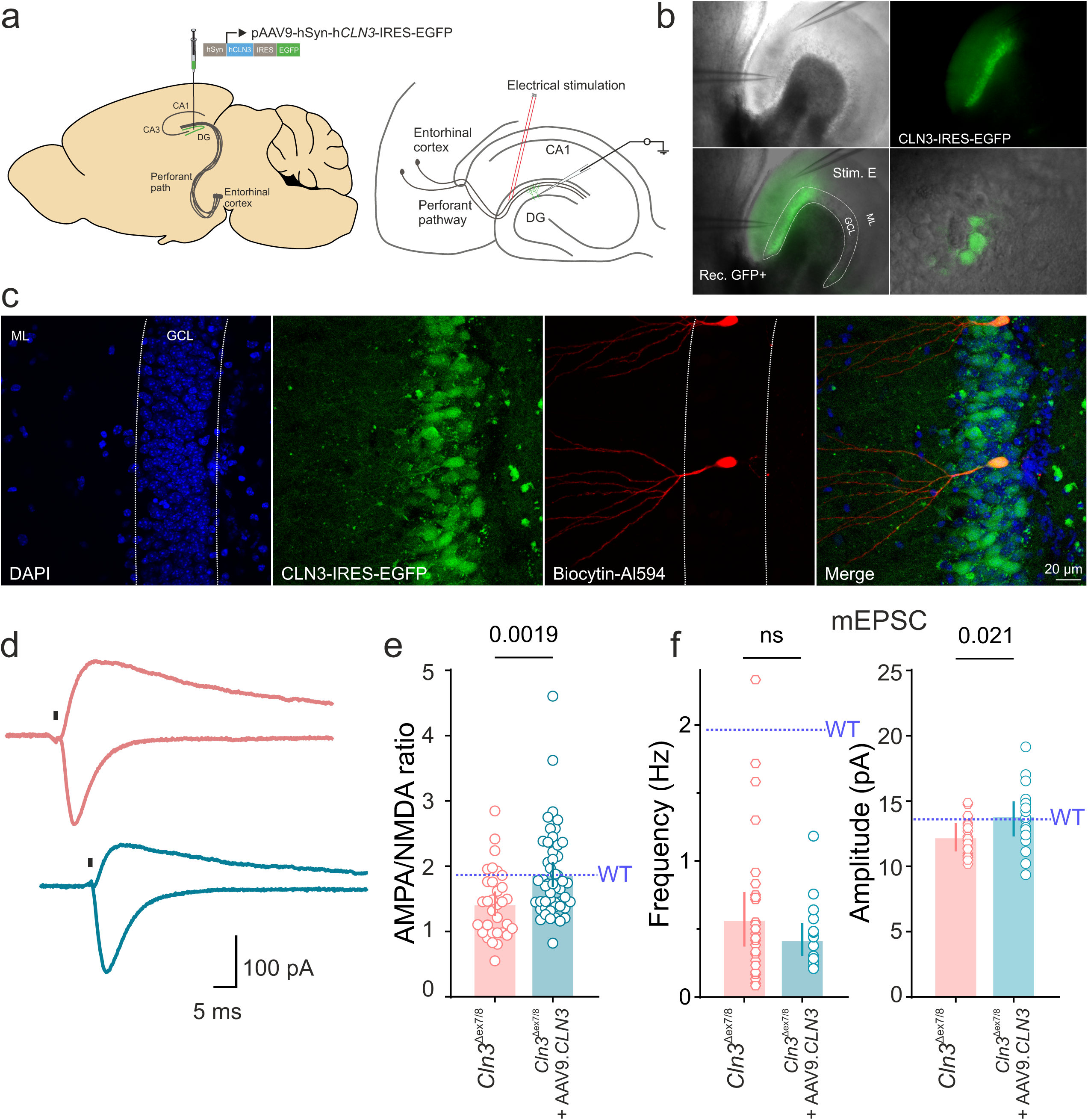
Postsynaptic re-expression of *CLN3* normalizes synaptic strength in *Cln3*^Δex7/8^ mice. (a) Schematic of AAV injections into the dentate gyrus to re-introduce *CLN3* in postsynaptic DG granule cells. (b) DIC-IR image showing CLN3-EGFP expression and positions of stimulation (Stim. E) and recording (rec. GFP+) electrodes during electrophysiological recordings. (c) Post hoc verification of patched cells by immunostaining after biocytin filling via patch pipette; merged image shows colocalization of CLN3-EGFP and biocytin, confirming successful targeting. (d) Example traces of evoked AMPA and NMDA receptor-mediated currents at holding potentials of -70 mV and +40 mV, respectively. Stimulus artifacts are removed for clarity (replaced with black vertical line). (e) Summary bar plots of AMPA/NMDA ratio (n= 33,45; N= 5,5 [*Cln3*^Δex7/8^, *Cln3*^Δex7/8^ + AAV9.*CLN3*]) and (f) mEPSC frequency and amplitude (n= 25, 22; N= 4,3 [*Cln3*^Δex7/8^, *Cln3*^Δex7/8^ + AAV9.*CLN3*]). Horizontal dashed line in blue above the data bars indicate WT values. Data are median [IQR] for AN ratio and mEPSC frequency and mean [95% CI] for mEPSC amplitude; ns, non-significant; p-values from Mann-Whitney tests and Unpaired t-test

## Discussion

Juvenile neuronal ceroid lipofuscinosis (JNCL) is a devastating disease that impairs neuronal function. Despite recent advances in understanding the molecular role of CLN3, it remains unclear how CLN3 deficiency leads to neuronal dysfunction and cognitive decline. To investigate the extent to which CLN3 loss directly affects neuronal performance, we analyzed intrinsic neuronal properties as well as pre- and postsynaptic function in young *Cln3*^Δex7/8^ mice. There is minimal storage burden and no neurodegeneration in *Cln3*-deficient mice at this early disease stage [2, 5, 10, 22, 35, 41, 42, 53, 54]. We found that neurons of *Cln3*^Δex7/8^ mice exhibit defects in intrinsic excitability, vesicle release probability, and synaptic strength. Furthermore, we could show that re-expression of *CLN3* after disease onset in either the presynaptic or postsynaptic compartment reversed these synaptic deficits and restored normal function. Together, our findings highlight the role of CLN3 in maintaining synaptic integrity and indicate the potential of targeted gene therapy to restore synaptic function in JNCL.

To screen for early onset synaptic deficits, we recorded mEPSCs and mIPSCs from dentate gyrus granule cells at 3 weeks and 4 months of age. *Cln3*^Δex7/8^ mice showed reduced frequency and amplitude of mEPSCs at these early-disease stages, whereas mIPSCs were unaffected. This clearly indicates that loss of CLN3 causes synaptic dysfunction independent of the occurrence of substantial storage accumulation and overt neuronal death [2, 5, 10, 22, 35, 41, 42, 53, 54]. The reduction of mEPSC frequency suggested impairments in presynaptic release while the reduced amplitude point to postsynaptic defects. This prompted us to conduct a more detailed analysis of pre- and postsynaptic function.

Presynaptically, we observed a reduction in synaptic vesicle release probability (P_r_) and an increased readily releasable pool (RRP) size, whereas recovery and replenishment rates of RRP remained unaffected. The altered P_r_ is consistent with impaired molecular assembly of the SNARE complex, a core component of the vesicle release machinery, as revealed by protein analyses in *Cln3*-deficient cortical neurons [43]. Although the precise mechanism underlying defective assembly of the release machinery remains unknown, it may reflect endolysosomal trafficking disturbances caused by CLN3 loss [6, 11, 44, 56]. Such perturbations can mis-sort synaptic vesicle (SV) cargo and favor formation of fusion-incompetent vesicles, thereby lowering P_r_ [16, 17]. Consistent with this, inhibition of endolysosomal trafficking in the presynapse have been shown to negatively regulate SV fusion by increasing the abundance of VAMP4 enriched SVs with reduced P_r_ [18]. The enlarged RRP phenotype may represent a compensatory response to reduced P_r_, aimed at stabilizing synaptic output by increasing the number of release-ready vesicles. Alternatively, impaired membrane trafficking in *Cln3*-deficient mice could mis-route vesicles between different presynaptic pools thus generating a larger pool with low fusion competence. This interpretation is supported by the interaction of CLN3 with the small GTPase Rab7a, a key regulator of endolysosomal cargo trafficking [56]. Additionally, proteomic profiling of synaptosome in *Cln3*^Δex7/8^ mice identified clusters of proteins associated with endocytosis and synaptic function with expression patterns matching regional vulnerability [33]. Collectively, these point to presynaptic endolysosomal dysfunction as a mechanism contributing to synaptic defects in *Cln3* disease.

The reduced mEPSC frequency could also be partly explained by the lower dendritic spine density, which reflects fewer excitatory synaptic contacts. This reduction in spine numbers may result from impaired autophagy. Several *Cln3-*deficient models report impairments in autophagic processes, including reduced autophagosome/lysosome fusion [6, 7, 34, 52]. Disruption of autophagy in hippocampal neurons have been shown to induce loss of dendritic spines and impair synaptic plasticity [9, 21]. Furthermore, impaired mTOR-autophagy signaling has been associated with defective spine pruning in postmortem brain samples from patients diagnosed with autism spectrum disorder [49]. Reduced dendritic spine density is often correlated with cognitive decline, suggesting that the observed decrease in dendritic spines may contribute to the cognitive deficits in JNCL. Moreover, we found that the spine loss was accompanied by subtle alterations in dendritic morphology suggesting an underlying neurodevelopmental component.

At the postsynapse, *Cln3*^Δex7/8^ neurons showed a reduced AMPA/NMDA ratio, indicating impaired AMPAR-mediated transmission and reduced excitatory drive. AMPARs are continuously shuttled between synaptic and extrasynaptic sites, with a fraction of receptors retrieved from the perisynaptic space through endocytic and autophagic processes [19, 36]. During plastic changes, this turnover is dramatically upregulated. Within the synapse, these receptors are sorted through the endolysosomal system, where they are either targeted for degradation, retained in recycling endosomes for later reinsertion, or shuttled back to the plasma membrane [39]. Given this tight coupling of synaptic activity and receptor turnover, disturbances in endolysosomal function, as observed in *Cln3*-deficient neurons [7, 34, 52], are expected to impair excitatory synaptic transmission. The weakening of excitatory drive observed here, together with the reduced release probability, may underlie the altered circuit excitability observed in our HD-MEA recordings and also reported by others [2, 46] and may help explain cognitive failure due to *Cln3*-deficiency.

The decrease in AMPAR-mediated transmission appears to contrast with earlier reports of increased AMPAR surface expression in *Cln3*^Δex1-6^ mice [27]. One possible explanation is that although synaptic AMPAR content is reduced, leading to impaired transmission, extrasynaptic expression is increased due to redistribution from synaptic sites. Such an imbalance could result from mis-trafficking of AMPARs through the endolysosomal system in *Cln3*-deficient neurons as proposed earlier [28]. At 3 weeks of age, we did not observe changes in mEPSC amplitude, consistent with previous reports of unaltered total AMPAR levels in *Cln3*^Δex1-6^ mice (P14-16) [48]. However, by 4 months we detected reduced mEPSC amplitude and A/N ratio suggesting a progressive nature of postsynaptic dysfunction.

Interestingly, inhibitory synapses appear to be affected later in the disease and to a lesser extent at early stages due to *Cln3* loss. At 14 months (symptomatic), we have previously demonstrated a loss of specific subclasses of GABAergic interneurons, which, together with synaptic dysfunction, results in circuit disinhibition [13]. However, at 3 weeks (presymptomatic), we found no change in inhibitory transmission, with only a slight trend toward reduced frequency at 4 months. This temporal difference, compared with prominent early excitatory defects, may represent distinct demands placed on the endolysosomal system by the glutamatergic vs GABAergic synapses. The preferential reduction in excitation, together with relatively preserved inhibition (E/I imbalance), suggests an inhibitory bias in the early network, which may explain the reduced hippocampal bursting and hypoexcitable phenotype observed in young *Cln3*- deficient mice without seizures. At late disease stages, when inhibition also deteriorates, network stability may be compromised, allowing seizures to emerge even though excitation remains weakened [2].

In addition to synaptic dysfunction, we observed aberrant intrinsic properties in dentate granule cells as early as 3 weeks of age. The overall reduction in excitability aligns with previously reported network and cellular deficits [8, 46] and may contribute to the hypoexcitable phenotype. By contrast, at 6 months of age, while DG granule cells remained hypoexcitable, we detected increased excitability in CA1 pyramidal neurons (Supplementary Fig. S3 and S4). This regional/cell type difference is in line with published observations and may explain the network excitability shift in the CA1 region [2]. The molecular mechanisms underlying these divergent responses between dentate granule cells and CA1 pyramidal neurons, however, remain to be elucidated [4].

Despite ongoing synaptic weakening, long-term potentiation (LTP), a classic cellular correlate of learning and memory, remained unaffected at the early disease stage, possibly reflecting homeostatic compensation. In contrast, spontaneous network activity recorded in acute hippocampal slices revealed a marked reduction in network bursting. This reduction can be explained by our findings at both the synaptic and neuronal level: reduced excitatory drive and impaired intrinsic excitability. Functional network bursts are essential for information encoding and retrieval in the dentate gyrus where they support pattern separation and coordinate oscillatory activity [32, 40]. Therefore, impairments in this critical network property provides a plausible mechanism for the cognitive deficits associated with *CLN3* mutations.

AAV9-mediated neuronal rescue of *CLN3* in 4-month-old *Cln3*^Δex7/8^ mice restored synaptic function, three weeks after viral injection, demonstrating the potential for postnatal therapeutic intervention. Targeted re-expression of *CLN3* in presynaptic and postsynaptic neurons improved vesicle release and synaptic strength, respectively, thereby not only halting functional decline but reversing it. This extends beyond earlier studies in which re-expression of *Cln3* in mice slowed disease progression when preformed at very early stages [1, 3, 20, 47] and supports the promise of AAV gene therapy as a treatment strategy in JNCL (NCT03770572; phase1/2 clinical trial in JNCL). At the mechanistic level, these results show that CLN3 is required at both synaptic sites to sustain function. They provide direct evidence that CLN3 maintains synaptic integrity and that synaptic deficits are not solely secondary to developmental disturbances or neurodegeneration. This is particularly significant since network alterations have been reported as early as the first postnatal week [46] and have been proposed to reflect a neurodevelopmental rather than purely neurodegenerative condition.

Functional synaptic recovery following AAV-mediated *CLN3* rescue raises new directions for investigation, including whether such recovery can translate into cognitive and behavioral improvements at later symptomatic stages. Our analyses focused on hippocampal principal cells and perforant pathway synapses, with subsequent work needed to determine whether CLN3 exerts neuronal-type or synapse-specific effects in other regions, such as thalamus or cerebellum, which also show early vulnerability in the disease. Future work extending these analyses across brain regions at early-disease stages will help define the full spectrum of the role of CLN3 in brain function and to optimize therapeutic strategies.

Impairments in synaptic function have been reported in other NCL subtypes, such as CLN1 and CLN10 [26, 31, 50], as well as in other lysosomal storage disorders, including MPSIIIC and NPC1 [38, 51, 55]. Evidence from some models indicates that synaptic dysfunction can emerge early in disease progression, yet the precise timing and underlying mechanisms remain incompletely understood. Distinguishing whether these alterations precede neurodegeneration or result from it requires direct functional assessment of neuronal and synaptic physiology across disease stages. Through detailed electrophysiological analyses, our study defines the onset and nature of synaptic dysfunction in CLN3 deficiency, thereby providing a functional framework for identifying early synaptic and neuronal impairments. These findings underscore the importance of establishing a temporal map of functional changes to determine the earliest and most effective therapeutic window.

In summary, we identify defective presynaptic vesicle release and reduced synaptic strength as early features of CLN3 deficiency, preceding overt pathology. These alterations, together with changes in intrinsic excitability, disrupt network activity. As synaptic signaling provides the basis for circuit-level processing, such deficits are likely to impair information encoding and integration, providing a mechanistic basis for the cognitive impairments observed in JNCL. Importantly, we demonstrate that postnatal, neuron-specific gene rescue restores synaptic function after disease onset. Together with earlier work showing benefits of neonatal interventions, our findings broaden the scope of AAV-based approaches by showing that reversal is also achievable at early symptomatic stages. In summary, our findings establish CLN3 as essential for synaptic integrity and highlight the potential of postnatal gene therapy.

## Author contributions

JvE and BG conceptualized, supervised and acquired funding for the study. MAW performed patch-clamp and optogenetics experiments and analyzed the data. MAW and CH performed HD-MEA recordings and analyzed the data. TM provided experimental advice related to HD-MEA recordings and supervised it. MAW and BG wrote the manuscript with critical feedback from all authors. All authors have reviewed, and approved the final manuscript.

## Acknowledgements

We would like to thank members of JvE laboratory and Dr. P. Herman van der Putten (NCL-Stiftung) for valuable discussions. Support from Dr. Petri Turunen and the IMB Microscopy Core Facility is gratefully acknowledged (DFG, project number 497669232).

