## Supplementary information for "Reversible synaptic deficits in early-stage Batten disease"

### Supplementray Figure S1

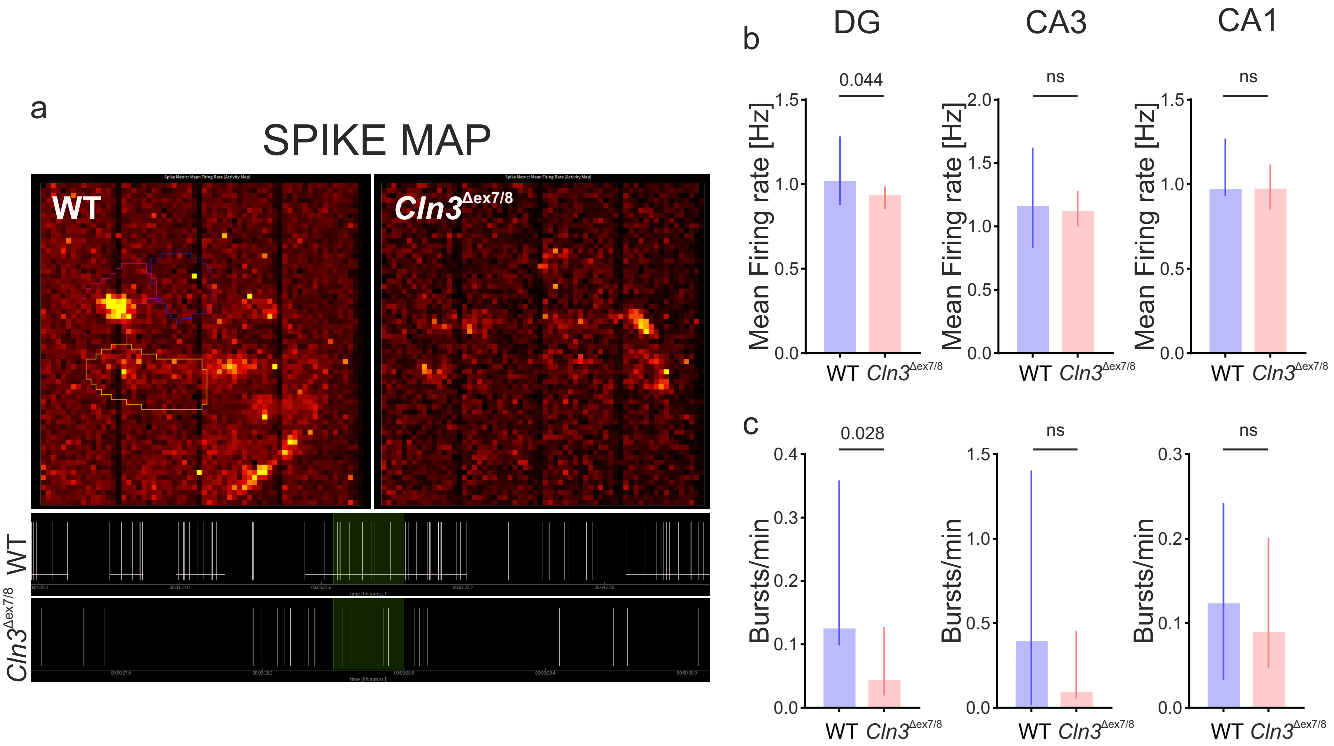

**Fig. S1** Reduced network bursting in the hippocampus of Cln3$^{\Delta ex7/8}$ mice. (a) Representative heat map of spontaneous activity (spikes) from acute hippocampal slices (4-month-old) recorded with HD-MEA. Bursts were defined as ≥5 consecutive spikes with inter-spike interval ≤100 ms (bottom panel: detected spikes grouped into bursts). (b) Summary bar plots of mean firing rate (MFR, Hz) and (c) bursting rate (bursts/min) in DG, CA3 and CA1 for WT and Cln3$^{\Delta ex7/8}$ slices (n= 10,14; N= 4,4 [WT, Cln3$^{\Delta ex7/8}$]). Data are mean [95% CI] for DG, CA3 MFR, and median [IQR] for all other bar plots; ns, non-significant; p-values from Unpaired t-tests and Mann-Whitney tests

### Supplemententray Figure S2

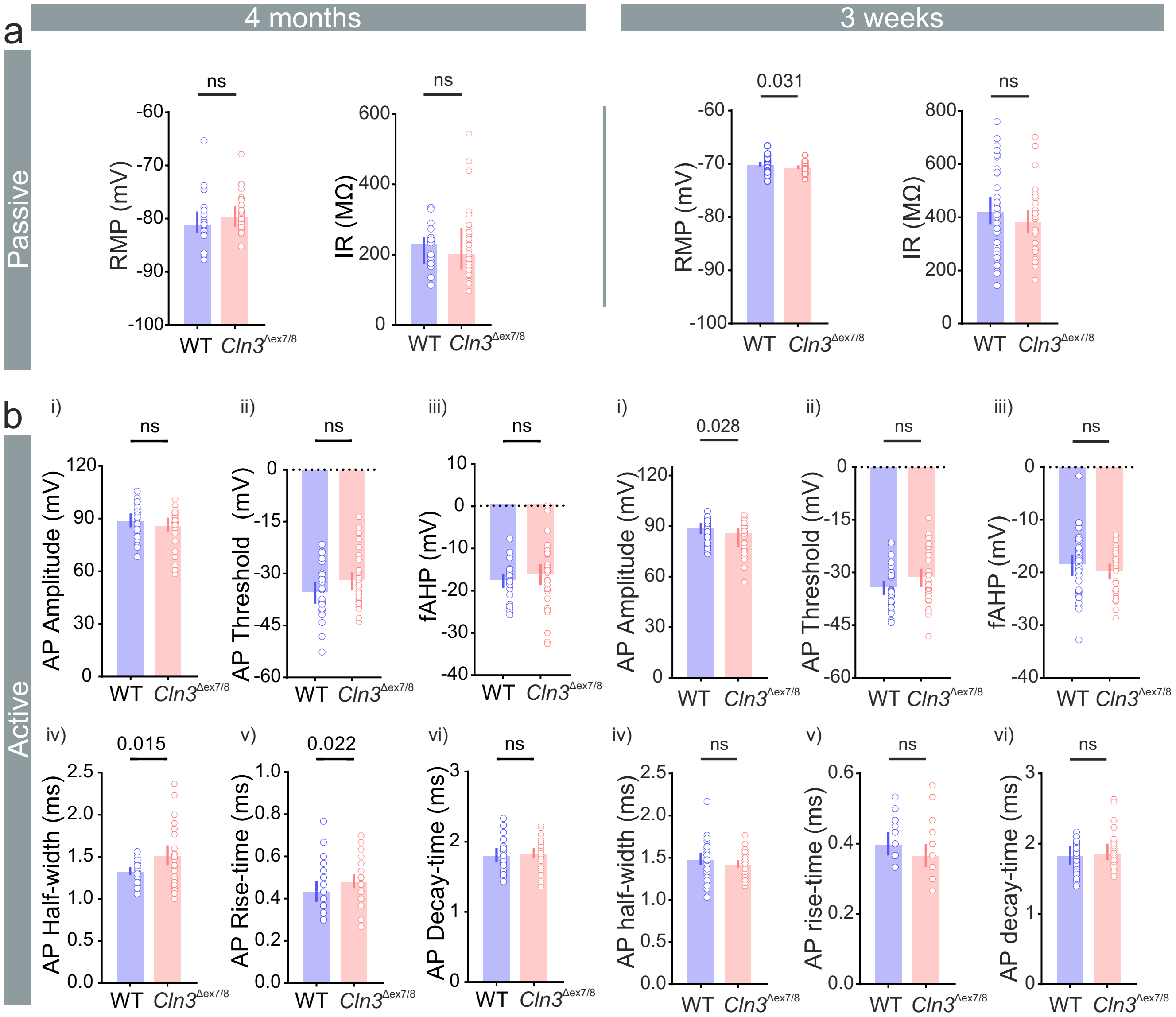

**Fig. S2** Intrinsic membrane properties of neurons in *Cln3*$^{\Delta ex7/8}$ mice. (a) Passive properties: summary bar plots of resting membrane potential (RMP) and input resistance (IR) at 4-months (left, median [IQR]; n= 20,32; N= 3,3 [WT, *Cln3*$^{\Delta ex7/8}$]] and at 3-weeks (right, mean [95% CI]; n= 38,33; N= 5,4 [WT, *Cln3*$^{\Delta ex7/8}$]]. (b) Active properties: summary bar plots of action potential (AP) amplitude, threshold, fast afterhyperpolarization (fAHP), half-width, rise-time, and decay-time at 4-months (left; n= 26,30; N= 4,5 [WT, *Cln3*$^{\Delta ex7/8}$]) and at 3-weeks (right; n= 27,24; N= 5,4 [WT, *Cln3*$^{\Delta ex7/8}$]). Data are mean [95% CI] for plots in 4-month (vi), 3-wks (ii, iii, iv), and median [IQR] for all other plots; ns, non-significant; p-values from Unpaired t-tests and Mann-Whitney tests

### Supplementray Figure S3

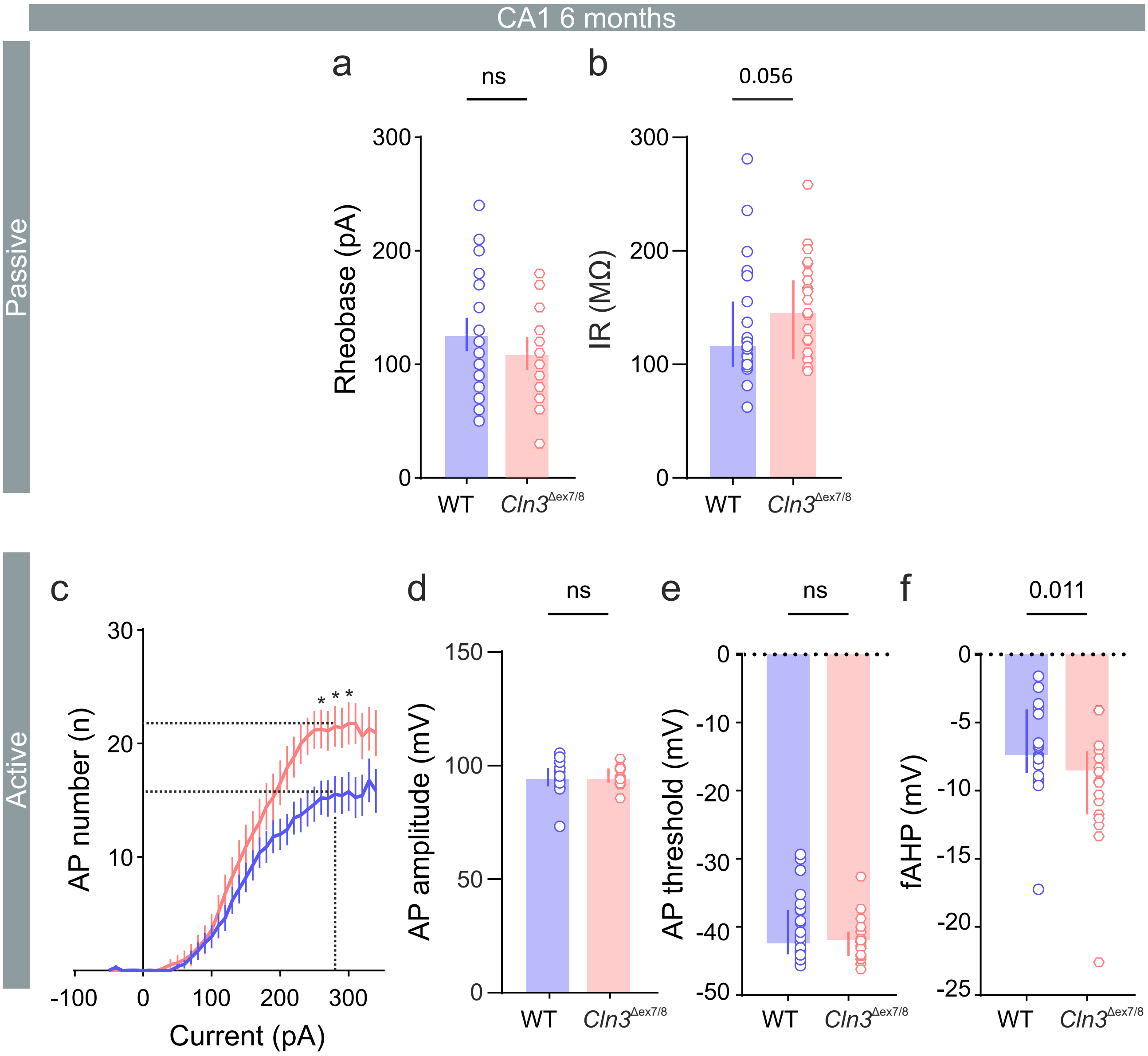

**Fig. S3** Intrinsic membrane properties of CA1 neurons at 6-months in *Cln3*$^{\Delta ex7/8}$ mice. (a, b) Summary bar plots of rheobase (pA) and input resistance (IR). (c) Number of action potentials (AP) plotted against increasing current steps, mean ± SEM. (d-f) Summary bar plots of AP amplitude, threshold, and fast afterhyperpolarization [n= 31,27; N= 5,5 [WT, *Cln3*$^{\Delta ex7/8}$]]. Data are mean ± SEM in (c) (*p < 0.05; Mann-Whitney tests), mean [95% CI] in (a) and median [IQR] (b, d-f); ns, non-significant; p-values from Unpaired t-test (a) and Mann-Whitney test (f)

### Supplementray Figure S4

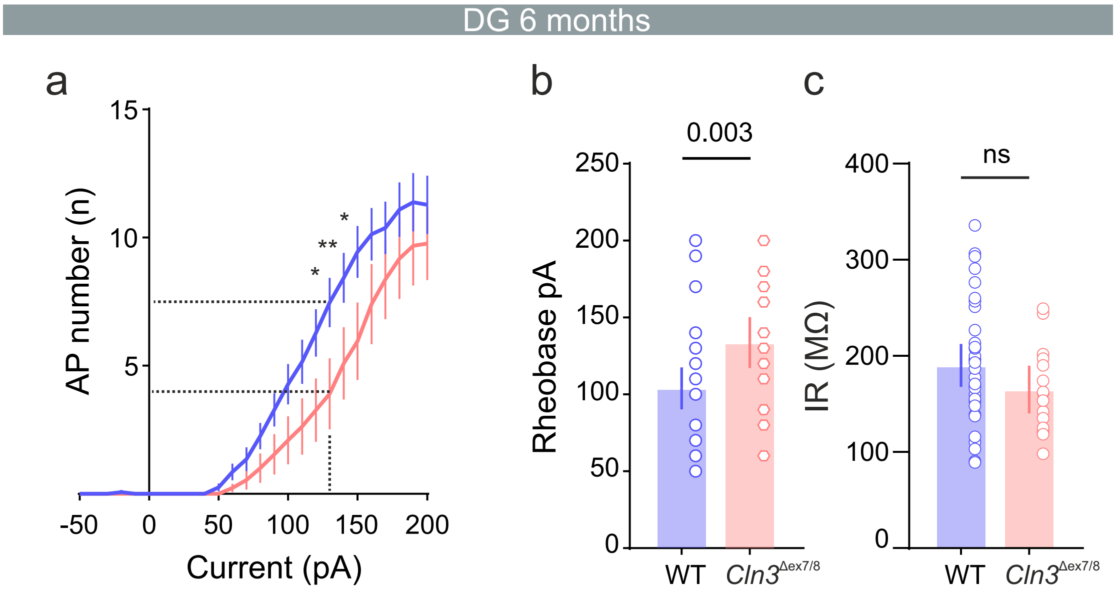

**Fig. S4** Intrinsic membrane properties of DG granule cells at 6-months in *Cln3*$^{\Delta ex7/8}$ mice. (a) Number of action potentials (AP) plotted against increasing current steps, mean ± SEM. (b) Summary bar plots of Rheobase (pA) and (c) input resistance (IR). Data are mean ± SEM in (a) (**$p < 0.01$, *$p < 0.05$; Mann-Whitney tests), median [IQR] in (b) and mean [95% CI] in (c); n= 33,21; N= 5,3 [WT, *Cln3*$^{\Delta ex7/8}$]); ns, non-significant; p-values from Unpaired t-test (IR) and Mann-Whitney test (rheobase)

### Supplementray Figure S5

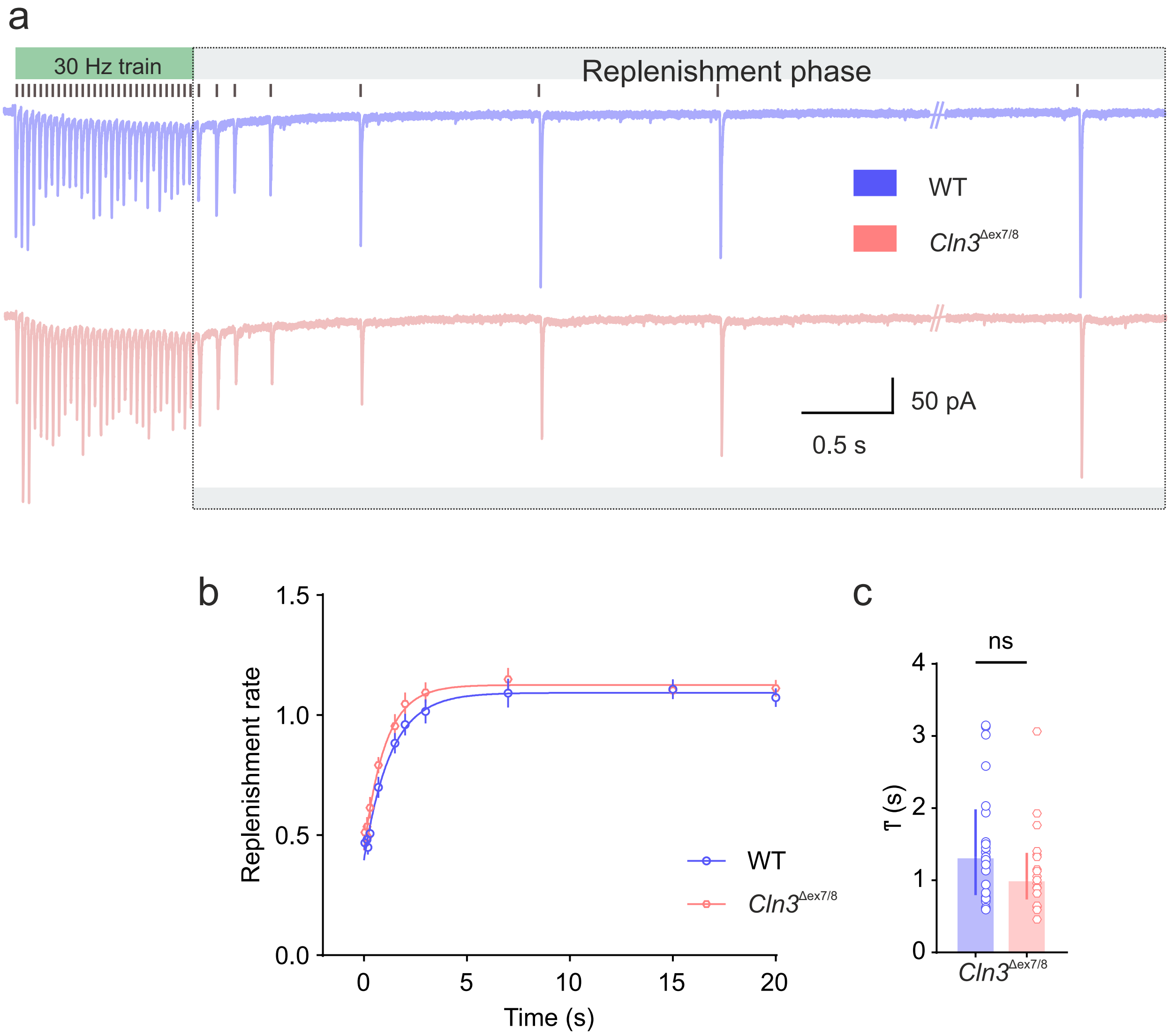

**Fig. S5** RRP replenishment rate after pool depletion is unaffected. (a) Representative trace of evoked EPSCs in response to 30 Hz perforant pathway stimulation, followed by single eEPSCs evoked at increasing inter-stimulus intervals (replenishment phase; dotted box, black lines above traces mark stimulus times). (b) Solid line represents single exponential decay fit of the data. (c) Summary bar plot of decay time constant (τ); n= 21,21; N= 4,3 [WT, *Cln3*$^{\Delta ex7/8}$]. Data are median [IQR]; ns, non-significant; Mann-Whitney test

### Supplementray Figure S6

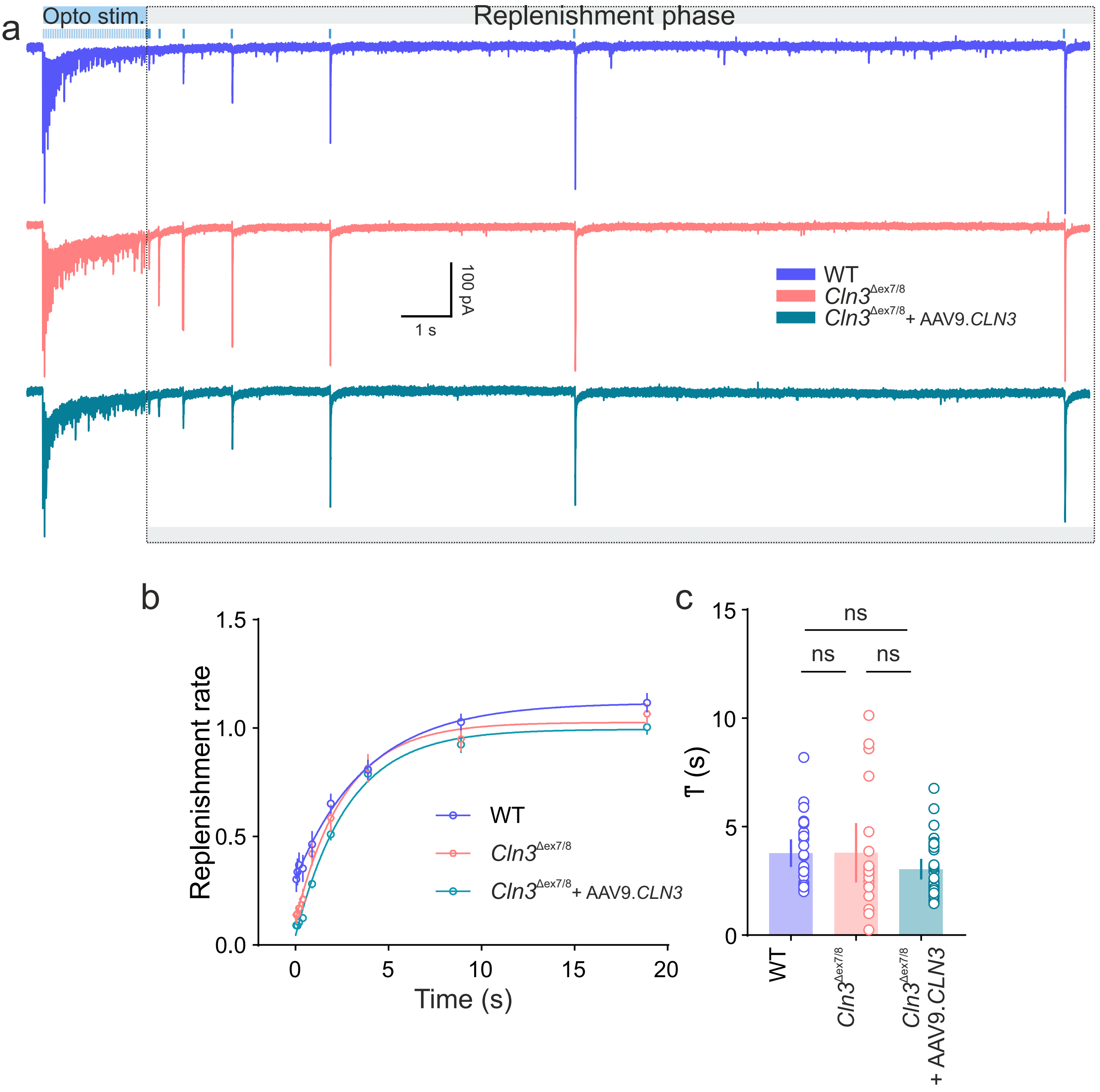

**Fig. S6** Presynaptic re-expression of *CLN3* in *Cln3*$^{\Delta ex7/8}$ mice does not affect RRP replenishment rate. (a) Representative trace of evoked EPSCs in response to 30 Hz optical stimulation of the perforant pathway, followed by single eEPSCs evoked at increasing inter-stimulus intervals (replenishment phase; dotted box, blue lines above the traces indicate stimulus times). (b) Solid line represents single exponential decay fit of the data. (c) Summary bar plot of decay time constant (τ); n= 22,20,26; N= 3,3,4 [WT, *Cln3*$^{\Delta ex7/8}$, *Cln3*$^{\Delta ex7/8}$ + AAV9.*CLN3*]. Data are median [IQR]; ns, non-significant; Kruskal-Wallis test followed by Dunn's multiple comparisons test

### Supplementray Figure S7

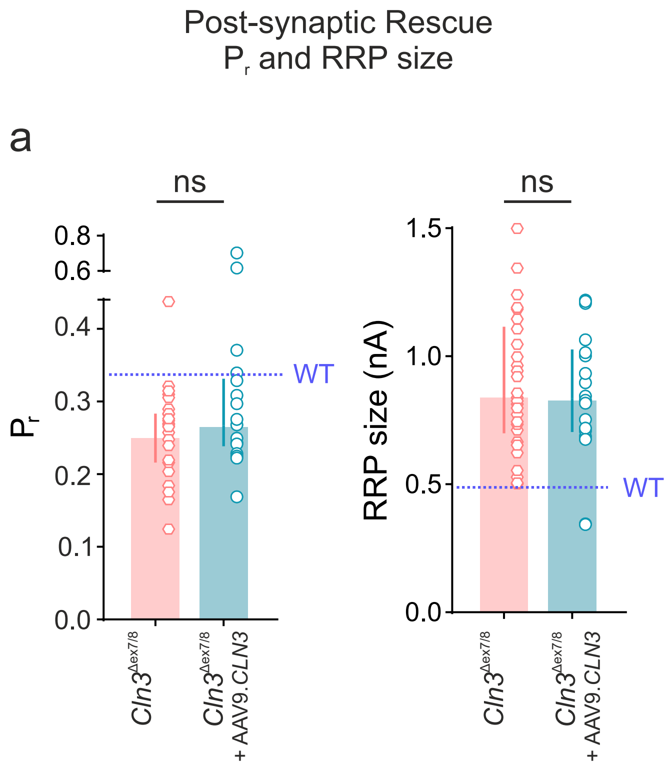

**Fig. S7** Postsynaptic re-expression of *CLN3* does not improve presynaptic function in *Cln3*$^{\Delta ex7/8}$ mice. (a) Summary bar plots of release probability ($P_r$), readily releasable pool (RRP) size (n= 28, 19; N= 4,3 [*Cln3*$^{\Delta ex7/8}$, *Cln3*$^{\Delta ex7/8}$ + AAV9.*CLN3*]). Data are median [IQR] for $P_r$ and mean [95% CI] for RRP size; ns, non-significant; Mann-Whitney test and Unpaired t test. Horizontal dashed line in blue above the data bars indicate WT values

Table 1. Intrinsic membrane properties of DG granule cells from WT and *Cln3*$^{ex7/8}$ mice at 3 weeks and 4 months of age

<table>
<tr><th colspan="2">Parameter</th><th>WT</th><th>Cln3$^{\Delta ex7/8}$</th><th>n & N (WT, Cln3$^{\Delta ex7/8}$)</th><th>Normality (Shapiro Wilk test)</th><th>Statistics</th></tr>
<tr><td rowspan="2">4 months</td><td>Input resistance (MΩ) median [P25 - P75]</td><td>233.80 [174.70 to 249.30]</td><td>204.00 [157.90 to 276.60]</td><td rowspan="2">n=20, 32 N=3,3</td><td>No</td><td>Mann Whitney test p=0.736</td></tr>
<tr><td>Resting membrane potential (mV) median [P25 - P75]</td><td>-80.91 [-82.73 to -78.64]</td><td>-79.53 [-81.56 to -77.54]</td><td>No</td><td>Mann Whitney test p=0.121</td></tr>
<tr><td rowspan="2">3 weeks</td><td>Input resistance (MΩ) mean [95% CI]</td><td>425.90 [374.40 to 477.40]</td><td>384.80 [341.80 to 427.70]</td><td rowspan="2">n=38, 33 N=5,4</td><td>Yes</td><td>Unpaired t test p=0.226</td></tr>
<tr><td>Resting membrane potential (mV) mean [95% CI]</td><td>-70.03 [-70.47 to -69.59]</td><td>-70.66 [-71.03 to -70.29]</td><td>Yes</td><td>Unpaired t test p=0.0314</td></tr>
</table>

<table>
<tr><th></th><th>Parameter</th><th colspan="3">Statistics</th><th>n & N (WT, Cln3$^{\Delta ex7/8}$)</th></tr>
<tr><td rowspan="4">4 months</td><td rowspan="4">Action Potential number vs current injection (pA)</td><td colspan="3">Multiple Mann Whitney test (Multiple comparisons FDR: Two-stage set-up (Benjamini, Krieger and Yakutieli))</td><td rowspan="4">n=26,30 N=4,5</td></tr>
<tr><td rowspan="3">Current step</td><td>160 pA</td><td>p=0.0100, q = 0.043</td></tr>
<tr><td>170 pA</td><td>p=0.0057, q = 0.043</td></tr>
<tr><td>180 pA</td><td>p=0.0117, q = 0.043</td></tr>
<tr><td rowspan="4">3 weeks</td><td rowspan="4">Action Potential number vs current injection (pA)</td><td colspan="3">Multiple Mann Whitney test (Multiple comparisons FDR: Two-stage set-up (Benjamini, Krieger and Yakutieli))</td><td rowspan="4">n=27,24 N=5,4</td></tr>
<tr><td rowspan="3">Current step</td><td>50 pA</td><td>p=0.0136, q = 0.0642</td></tr>
<tr><td>60 pA</td><td>p=0.0063, q = 0.0632</td></tr>
<tr><td>70 pA</td><td>p=0.0169, q = 0.0642</td></tr>
</table>

| Action potential (AP) Kinetics (4-month-old) | | | | | |
|---|---|---|---|---|---|
| | Parameter | WT | *Cln3*$^{\Delta ex7/8}$ | Normality (Shapiro Wilk test) | Statistics n=26,30; N=4,5 |
| 4 months | AP threshold voltage (mV) median [P25 - P75] | -37.54 [-40.57 to -30.15] | -33.68 [-37.97 to -29.16] | No | Mann Whitney test p=0.126 |
| | AP Amplitude (mV) median [P25 - P75] | 91.50 [82.21 to 95.78] | 88.13 [84.81 to 92.02] | No | Mann Whitney test p=0.263 |
| | AP Rise time (ms) median [P25 - P75] | 0.37 [0.37 to 0.51] | 0.50 [0.40 to 0.53] | No | Mann Whitney test p=0.0221 |
| | AP Decay time (ms) mean [95% CI] | 1.81 [1.72 to 1.91] | 1.84 [1.77 to 1.91] | Yes | Unpaired t test p=0.656 |
| | fast Afterhyperpolarization (fAHP) (mV) median [P25 - P75] | -17.38 [-19.87 to -15.29] | -15.59 [-19.82 to -12.57] | No | Mann Whitney test p=0.293 |
| | AP Half-width (ms) median [P25 - P75] | 1.35 [1.23 to 1.41] | 1.50 [1.24 to 1.73] | No | Mann Whitney test p=0.0151 |

| Action potential (AP) Kinetics (3-week-old) | | | | | |
|---|---|---|---|---|---|
| | Parameter | WT | *Cln3*$^{\Delta ex7/8}$ | Normality (Shapiro Wilk test) | Statistics n=27,24; N=5,4 |
| 3 weeks | AP threshold voltage (mV) median [P25 - P75] | -34.40 [-36.44 to -32.35] | -31.51 [-34.16 to -28.86] | Yes | Unpaired t test p=0.0841 |
| | AP Amplitude (mV) median [P25 - P75] | 89.12 [85.31 to 91.79] | 86.52 [77.86 to 88.90] | No | Mann Whitney test p=0.0289 |
| | AP Rise time (ms) median [P25 - P75] | 0.40 [0.37 to 0.43] | 0.37 [0.33 to 0.40] | No | Mann Whitney test p=0.389 |
| | AP Decay time (ms) mean [95% CI] | 1.83 [1.70 to 1.97] | 1.87 [1.77 to 2.00] | No | Mann Whitney test p=0.285 |
| | fast Afterhyperpolarization (fAHP) (mV) median [P25 - P75] | -18.61 [-20.61 to -16.60] | -19.82 [-21.32 to -18.32] | Yes | Unpaired t test p=0.329 |
| | AP Half-width (ms) median [P25 - P75] | 1.49 [1.42 to 1.56] | 1.43 [1.38 to 1.47] | Yes | Unpaired t test p=0.157 |

Table 2. Synaptic transmission: mEPSC and mIPSC values of DG granule cells from WT and *Cln3*$^{ex7/8}$ mice at 3 weeks and 4 months of age

| Parameter | | | WT | *Cln3*$^{\Delta ex7/8}$ | n & N (WT, *Cln3*$^{\Delta ex7/8}$) | Normality (Shapiro Wilk test) | Statistics |
|---|---|---|---|---|---|---|---|
| mEPSC | 4 months | Frequency (Hz)<br>median [P25 - P75] | 1.96<br>[1.61 to 2.79] | 0.75<br>[0.53 to 1.28] | n= 17,26;<br>N=3,3 | No | Mann Whitney test<br>p<0.0001 |
| | | Amplitude (pA)<br>mean [95% CI] | 13.61<br>[12.45 to 14.76] | 11.41<br>[10.48 to 12.34] | | Yes | Unpaired t test<br>p=0.0037 |
| | 3 weeks | Frequency (Hz)<br>median [P25 - P75] | 3.16<br>[2.27 to 4.61] | 2.11<br>[1.55 to 2.66] | n= 28,31;<br>N= 3,3 | No | Mann Whitney test<br>p=0.0048 |
| | | Amplitude (pA)<br>median [P25 - P75] | 15.82<br>[13.57 to 17.36] | 15.56<br>[13.90 to 16.84] | | No | Mann Whitney test<br>p=0.769 |

| Parameter | | | WT | *Cln3*$^{\Delta ex7/8}$ | n & N (WT, *Cln3*$^{\Delta ex7/8}$) | Normality (Shapiro Wilk test) | Statistics |
|---|---|---|---|---|---|---|---|
| mIPSC | 4 months | Frequency (Hz)<br>median [P25 - P75] | 2.61<br>[1.82 to 3.43] | 2.04<br>[1.48 to 2.74] | n= 31,40;<br>N=5,5 | No | Mann Whitney test<br>p=0.0784 |
| | | Amplitude (pA)<br>median [P25 - P75] | 32.01<br>[24.90 to 37.59] | 27.96<br>[21.48 to 33.58] | | No | Mann Whitney test<br>p=0.0588 |
| | 3 weeks | Frequency (Hz)<br>mean [95% CI] | 2.75<br>[1.20 to 4.30] | 1.80<br>[1.08 to 2.51] | n= 12,13;<br>N= 2,3 | Yes | Unpaired t test<br>p=0.206 |
| | | Amplitude (pA)<br>median [P25 - P75] | 21.12<br>[17.82 to 43.71] | 24.97<br>[20.08 to 36.54] | | No | Mann Whitney test<br>p=0.689 |

Table 3. AMPA/NMDA ratio values of DG granule cells from 4-month-old WT and *Cln3*$^{ex7/8}$ mice

| Parameter | WT | *Cln3*$^{\Delta ex7/8}$ | n & N (WT, *Cln3*$^{\Delta ex7/8}$) | Normality (Shapiro Wilk test) | Statistics |
|---|---|---|---|---|---|
| AMPA-NMDA ratio mean [95% CI] | 1.86 [1.65 to 2.08] | 1.18 [1.01 to 1.35] | n= 33,42; N=5,6 | Yes | Unpaired t test p <0.0001 |
| NMDAR decay τ (ms) mean [95% CI]] | 43.38 [37.50 to 49.26] | 44.61 [38.99 to 50.22] | n= 22,30; N=2,2 | Yes | Unpaired t test p= 0.762 |

Table 4. Dendritic spine density of DG granule cells from 4-month-old WT and *Cln3*$^{ex7/8}$ mice

| Parameter | WT | *Cln3*$^{\Delta ex7/8}$ | n & N (WT, *Cln3*$^{\Delta ex7/8}$) | Normality (Shapiro Wilk test) | Statistics |
|---|---|---|---|---|---|
| Spine count (per 10 μm) mean [95% CI] | 18.8 [17.9 to 19.7] | 16.05 [14.6 to 17.4] | n= 43,39; N=8,5 | Yes | Unpaired t test p=0.0008 |
| Spine count (per 10 μm) subtype - Mushroom mean [95% CI] | 1.83 [1.38 to 2.67] | 1.28 [0.74 to 2.12] | | No | Mann Whitney test<br>Multiple comparisons FDR<br>Two-stage set-up (Benjamini, Krieger and Yakutieli)<br><br>Mushroom: p= 0.01163 q=0.025<br>Thin: p= 0.0167 q=0.025<br>Stubby: p= 0.252 q=0.255 |
| Spine count (per 10 μm) subtype - Thin mean [95% CI] | 11.61 [10.09 to 13.49] | 9.70 [7.34 to 13.2] | | No | |
| Spine count (per 10 μm) subtype - Stubby mean [95% CI] | 4.53 [3.59 to 5.53] | 4.31 [3.55 to 5.21] | | No | |
| Total dendritic length (mm) median [P25 - P75] | 43.38 [37.50 to 49.26] | 44.61 [38.99 to 50.22] | n= 17,26; N=3,5 | No | Mann Whitney test p= 0.877 |

Table 5. Readily-releasable pool (RRP) values of DG granule cells from 4-month-old WT and *Cln3*$^{ex7/8}$ mice

| Parameter | WT | *Cln3*$^{\Delta ex7/8}$ | n & N (WT, *Cln3*$^{\Delta ex7/8}$) | Normality (Shapiro Wilk test) | Statistics |
|---|---|---|---|---|---|
| RRP size (nA)<br>median [P25 - P75] | 0.629<br>[0.498 to 0.782] | 0.815<br>[0.62 to 1.02] | n= 31,25;<br>N=4,3 | No | Mann Whitney test<br>p=0.003 |
| $P_r$ (release probability)<br>mean [95% CI] | 0.34<br>[0.31 to 0.38] | 0.27<br>[0.24 to 0.31] | | Yes | Unpaired t test<br>p= 0.0038 |
| Slope<br>median [P25 - P75] | 0.10<br>[0.08 to 0.13] | 0.10<br>[0.08 to 0.14] | | No | Mann Whitney test<br>p= 0.662 |
| Replenishment rate τ (ms)<br>median [P25 - P75] | 1.32<br>[0.79 to 1.98] | 1.007<br>[0.733 to 1.38] | | No | Mann Whitney test<br>p= 0.132 |

Table 6. Hippocampal LTP in 4-month-old WT and *Cln3*$^{ex7/8}$ mice

| | Parameter | WT | *Cln3*$^{\Delta ex7/8}$ | n & N (WT, *Cln3*$^{\Delta ex7/8}$) | Statistics |
|---|---|---|---|---|---|
| Induced | First 10 min<br>EPSC peak %<br>mean [95% CI] | 207.9<br>[162.1 to 253.8] | 183.4<br>[138.0 to 228.8] | n= 13,14;<br>N=6,5 | 2-way RM ANOVA with Sidak's post hoc test<br>F (1, 25) = 0.115 p=0.736 (Genotype) |
| | Last 10 min<br>EPSC peak %<br>mean [95% CI] | 181.1<br>[140.3 to 221.9] | 183.8<br>[129.6 to 238.0] | | |
| Control | First 10 min<br>EPSC peak %<br>mean [95% CI] | 68.68<br>[57.50 to 79.86] | 64.99<br>[52.14 to 77.83] | | 2-way RM ANOVA with Sidak's post hoc test<br>F (1, 25) = 0.267 p=0.610 (Genotype) |
| | Last 10 min<br>EPSC peak %<br>mean [95% CI] | 72.13<br>[54.77 to 89.48] | 67.08<br>[54.87 to 79.29] | | |

Table 7. Presynaptic properties recorded from DG-GCs after AAV9-mediated *CLN3* re-expression in presynaptic perforant pathway projections from Entorhinal cortex to DG in *Cln3*$^{ex7/8}$ mice

| Parameter | WT | *Cln3*$^{\Delta ex7/8}$ | *Cln3*$^{\Delta ex7/8}$ + AAV9.*CLN3* | n & N (WT, *Cln3*, *Cln3*$^{\Delta ex7/8}$ + AAV9.*CLN3)* | Normality (Shapiro Wilk test) | Statistics Kruskal-Wallis test followed by Dunn's multiple comparisons test |
|---|---|---|---|---|---|---|
| RRP size (nA) median [P25 - P75] | 0.477 [0.352 to 0.702] | 0.920 [0.598 to 1.12] | 0.632 [0.497 to 0.770] | n= 22,20,26; N=3,3,4 | No | WT vs *Cln3*$^{\Delta ex7/8}$ p=0.0003<br>WT vs *Cln3*$^{\Delta ex7/8}$ + AAV9.*CLN3* p=0.435<br>*Cln3*$^{\Delta ex7/8}$ vs *Cln3*$^{\Delta ex7/8}$ + AAV9.*CLN3* p=0.024 |
| $P_r$ (release probability) median [P25 - P75] | 0.277 [0.240 to 0.396] | 0.186 [0.136 to 0.262] | 0.257 [0.233 to 0.338] | | No | WT vs *Cln3*$^{\Delta ex7/8}$ p=0.0004<br>WT vs *Cln3*$^{\Delta ex7/8}$ + AAV9.*CLN3* p=0.904<br>*Cln3*$^{\Delta ex7/8}$ vs *Cln3*$^{\Delta ex7/8}$ + AAV9.*CLN3* p=0.008 |
| Slope median [P25 - P75] | 0.013 [0.009 to 0.018] | 0.012 [0.009 to 0.017] | 0.010 [0.007 to 0.014] | | No | WT vs *Cln3*$^{\Delta ex7/8}$ p>0.999<br>WT vs *Cln3*$^{\Delta ex7/8}$ + AAV9.*CLN3* p=0.433<br>*Cln3*$^{\Delta ex7/8}$ vs *Cln3*$^{\Delta ex7/8}$ + AAV9.*CLN3* p=0.384 |
| Replenishment rate τ (ms) median [P25 - P75] | 3.41 [2.38 to 4.84] | 2.85 [2.21 to 4.76] | 2.44 [1.97 to 4.17] | | No | WT vs *Cln3*$^{\Delta ex7/8}$ p=0.972<br>WT vs *Cln3*$^{\Delta ex7/8}$ + AAV9.*CLN3* p=0.124<br>*Cln3*$^{\Delta ex7/8}$ vs *Cln3*$^{\Delta ex7/8}$ + AAV9.*CLN3* p>0.999 |

Table 8. Pre- and postsynaptic properties recorded from DG-GCs after AAV9-mediated *CLN3* re-expression in postsynaptic granule cells of DG in *Cln3*$^{ex7/8}$ mice

| Parameter | *Cln3*$^{\Delta ex7/8}$ | *Cln3*$^{\Delta ex7/8}$ + AAV9.*CLN3* | n & N (*Cln3*, *Cln3*$^{\Delta ex7/8}$ + AAV9.*CLN3)* | Normality (Shapiro Wilk test) | Statistics |
|---|---|---|---|---|---|
| AMPA-NMDA ratio median [P25 - P75] | 1.43 [0.98 to 1.76] | 1.73 [1.39 to 2.33] | n= 33,45; N=5,5 | No | Mann Whitney test p= 0.0019 |

| | Parameter | *Cln3*$^{\Delta ex7/8}$ | *Cln3*$^{\Delta ex7/8}$ + AAV9.*CLN3* | n & N (*Cln3*, *Cln3*$^{\Delta ex7/8}$ + AAV9.*CLN3)* | Normality (Shapiro Wilk test) | Statistics |
|---|---|---|---|---|---|---|
| Presynaptic metrics after postsynaptic rescue | mEPSC Frequency (Hz) median [P25 - P75] | 0.40 [0.179 to 0.738] | 0.33 [0.267 to 0.506] | n= 25,22; N=4,3 | No | Mann Whitney test p= 0.739 |
| | mEPSC Amplitude (pA) mean [95% CI] | 12.42 [11.84 to 13.01] | 13.82 [12.62 to 15.01] | | Yes | Unpaired t test p= 0.021 |
| | RRP size (nA) mean [95% CI] | 0.893 [0.797 to 0.989] | 0.849 [0.721 to 0.976] | n= 28,19; N=4,3 | Yes | Unpaired t test p= 0.565 |
| | $P_r$ (release probability) median [P25 - P75] | 0.253 [0.216 to 0.283] | 0.268 [0.238 to 0.316] | | No | Mann Whitney test p= 0.147 |
